# Immune Selection Pressure Contributes to Pancreatic Cancer Immune Evasion

**DOI:** 10.1101/2020.06.15.151274

**Authors:** Reham Ajina, Annie Zuo, Shangzi Wang, Maha Moussa, Connor J. Cooper, Yue Shen, Quentin R. Johnson, Jerry M. Parks, Jeremy C. Smith, Marta Catalfamo, Elana J. Fertig, Sandra A. Jablonski, Louis M. Weiner

**Affiliations:** Department of Oncology and Lombardi Comprehensive Cancer Center, Georgetown University Medical Center, Washington, DC, USA; Department of Microbiology and Immunology, Georgetown University Medical Center, Washington, DC, USA; Graduate School of Genome Science and Technology, University of Tennessee, Knoxville, Tennessee, USA; UT/ORNL Center for Molecular Biophysics, Biosciences Division, Oak Ridge National Laboratory, Oak Ridge, Tennessee, USA; Department of Biochemistry and Cellular and Molecular Biology, University of Tennessee, Knoxville, Tennessee, USA; Department of Oncology, Sidney Kimmel Comprehensive Cancer Center, Johns Hopkins University School of Medicine, Baltimore, Maryland, USA; Department of Applied Mathematics & Statistics, Johns Hopkins University Whiting School of Engineering, Baltimore, Maryland, USA; Department of Biomedical Engineering, Johns Hopkins University School of Medicine, Baltimore, Maryland, USA

## Abstract

Pancreatic ductal adenocarcinoma (PDAC) is the third leading cause of cancer death in the United States. Pancreatic tumors are minimally infiltrated by T cells and are largely refractory to immunotherapy. Accordingly, the role of T cell immunity in pancreatic cancer has been somewhat overlooked. Here, we hypothesized that immune evasion in pancreatic cancer is induced in response to T cell-based immune selection pressure, and that understanding how pancreatic tumors respond to immune attack may facilitate the development of more effective therapeutic strategies. We now provide the first evidence that T cell-dependent host immune responses induce a PDAC-derived myeloid mimicry phenomenon and stimulate immune evasion. mT3-2D cells derived from a *Kras*^+/*LSL-G12D*^; *Trp53*^+/*LSL-R172H*^; *Pdx1-Cre* (KPC) mouse model of pancreatic cancer were grown in immunocompetent and immunodeficient C57BL/6 mice, and analyzed to determine the impacts of adaptive immunity specifically on malignant epithelial cells as well as on whole tumors. We found that immune selection pressure, via signal transducer and activator of transcription 1 (STAT1), stimulates malignant epithelial pancreatic cells to induce the expression of genes typically expressed by myeloid cells and alters intratumoral immunosuppressive myeloid cell profiles. Targeting the Janus Kinase (JAK)/STAT signaling pathway using the FDA approved drug, ruxolitinib, overcomes these tumor-protective responses and improves anti-PD1 antibody therapeutic efficacy. These findings provide future directions for treatments that specifically disable this mechanism of resistance in PDAC.

## Introduction

Worldwide, nearly half a million people are diagnosed with pancreatic cancer every year (1), and the mortality of these patients within 5 years is more than 90% (2). Therefore, there is an urgent need to develop more effective therapeutic strategies.

Pancreatic tumors typically have minimal infiltration of T cells, and thus pancreatic cancer is considered to be an immunologically cold neoplasm. Also, when T cells infiltrate pancreatic tumors, they are generally skewed toward T cells that exhibit pro-tumor properties rather than anti-tumor cytotoxic T cells (3). Even when effective cytotoxic cells successfully infiltrate pancreatic tumors, they are likely to either get suppressed or physically trapped within the stroma (4). Moreover, the pancreatic cancer myelosuppressive microenvironment has been shown to be derived by tumor-intrinsic properties (5,6). Although these observations could suggest that T cell immunity is not relevant to pancreatic cancer and that the pancreas has organ site-specific resistance to immunotherapy, recent studies have shown otherwise. Indeed, tumors of rare long-term pancreatic cancer survivors elicit more potent cytotoxic T cell responses compared to the rest of pancreatic cancer patients (7). Also, immune checkpoint blockades, which promote T cell activity, have demonstrated efficacy in ~2% of pancreatic cancer patients that exhibit high microsatellite instability (MSI-high) (8). These findings suggest that when the T cell anti-tumor immune response is sufficiently induced, T cells become effective weapons.

Many therapeutic interventions have been developed to target known immune escape routes in pancreatic cancer. However, the vast majority of pancreatic cancer immunotherapy clinical trials have failed (9). Hence, further resolution of the complex biology of pancreatic cancer immunity is needed.

Here, we hypothesized that immune evasion in pancreatic cancer is induced in response to immune selection pressure, and that understanding how pancreatic tumors respond to immune attack may facilitate the development of more effective therapeutic strategies. To test this hypothesis, mT3-2D pancreatic tumors were grown in immunocompetent wild-type (WT) and severe combined immunodeficiency (SCID) C57BL/6 mice. These tumors were then analyzed to determine the impacts of adaptive immunity on malignant epithelial cells and whole tumors. First, we determined that despite the induction of T cell immunity in immunocompetent mice bearing mT3-2D tumors, T cells failed to completely regulate tumor growth. Subsequently, we showed that immune selection pressure, potentially via JAK/STAT signaling, induced the expression of myeloid-associated genes in malignant epithelial cells and stimulated the infiltration of PD-L1-expressing myeloid cells. Targeting the JAK/STAT signaling pathway using the FDA approved drug ruxolitinib inhibited the expression of myeloid cell-associated genes and improved anti-PD1 therapeutic efficacy in mice. Together, these data suggest that some known features of the pancreatic cancer immunosuppressive microenvironment are driven in response to immune selection pressure. Disabling such responses offers new approaches to pancreatic cancer-directed immunotherapy.

## Results

### T cell Immunity Incompletely Regulates Pancreatic Cancer Tumor Growth in Syngeneic Murine Models

To determine if T cell immunity can control pancreatic tumor growth, mT3-2D cells were inoculated into syngeneic WT and SCID C57BL/6 mice. mT3-2D tumors in SCID mice grew more rapidly than did mT3-2D tumors in WT mice (**Figure 1a** **and Supplementary Figure 1A)**. A similar tumor growth pattern was observed in the PANC02 model, which is not characterized by mutations in *kras or p53* **(Supplementary Figure 1B-C)**. Using hematoxylin and eosin (H&E), Masson’s trichrome, and Ki67 IHC staining, there was no apparent difference in the histology, the level of fibrosis or the cellular proliferation rate between mT3-2D WT and SCID tumors, respectively **(Supplementary Figure 2)**.

**Figure 1.**
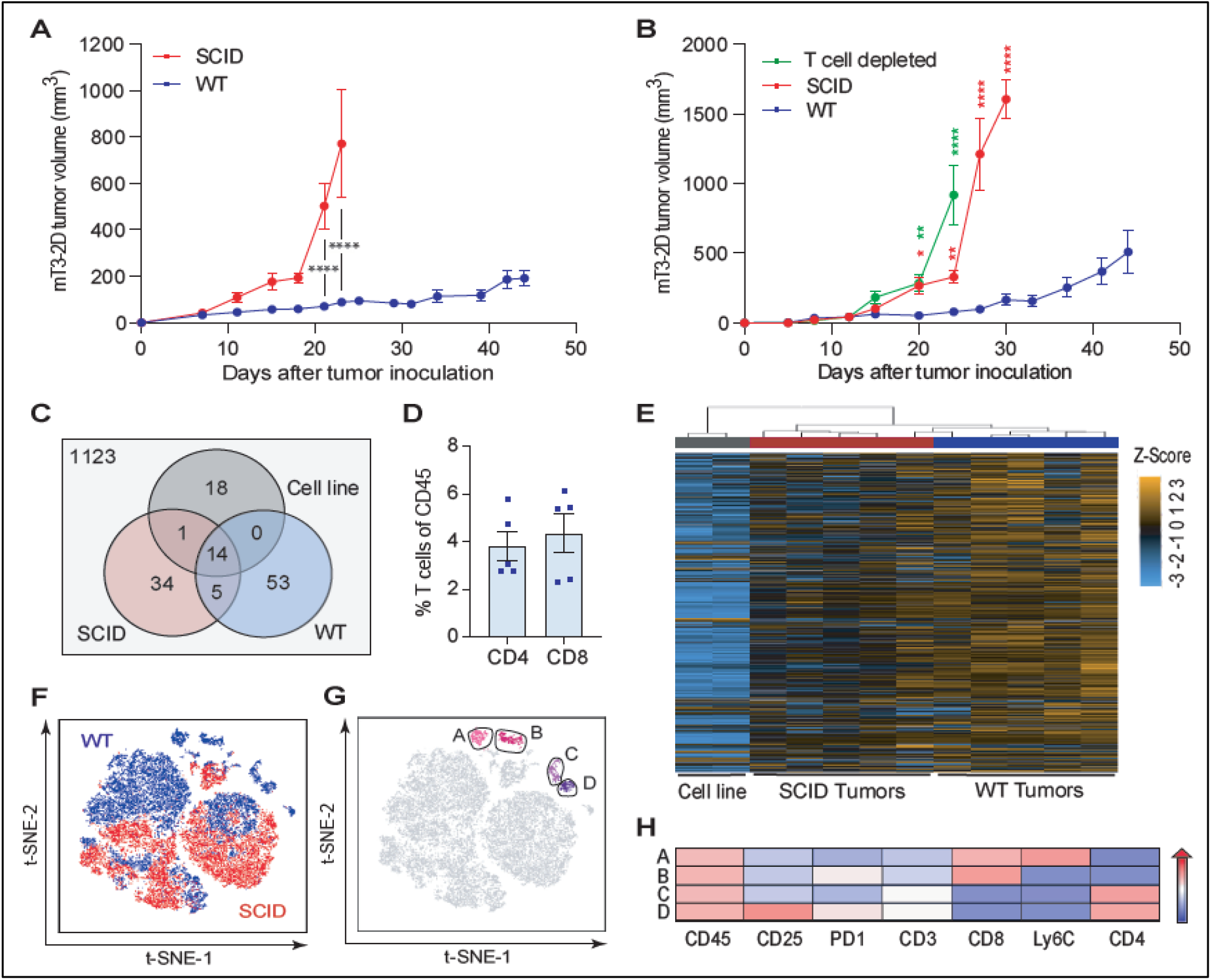
T cell immunity incompletely regulates mT3-2D tumor growth. **(A)** Growth curves depicting average subcutaneous mT3-2D tumor volumes (±SEM) grown in syngeneic WT (n=9) and SCID (n=13) C57BL/6J mice. A two-way ANOVA followed by Bonferroni multiple comparison test was used for the analysis. Experiment was repeated twice. **(B)** Growth curves depicting average subcutaneous mT3-2D tumor volumes (±SEM) grown in WT, SCID and T cell depleted C57BL/6J mice (n=10 per group). A two-way ANOVA followed by Tukey multiple comparison test was used for the analysis up to the time point “day 24”. A two-way ANOVA followed by Bonferroni multiple comparison test was used for the analysis of “day 27” and “day 30” time points. Green asterisks indicate statistical significance between WT and T cell depleted tumors, and red asterisks indicate statistical significance between WT and SCID tumors. Experiment was repeated twice. **(C)** Numbers of neoepitopes predicted by pVACtools for the mT3-2D cells, combined 5 SCID tumors, and combined 5 WT tumors. **(D)** Percentage of T cells (CD3^+^ CD4^+^ CD8^−^ or CD3^+^ CD4^−^ CD8^+^ cells) in immunocompetent tumors detected by flow cytometry and represented as frequency of CD45^+^ cells using FlowJo (n=5). **(E)** Unsupervised hierarchical clustering of normalized differentially expressed genes in the mT3-2D cell line (n=2 technical replicates, labeled with a gray bar at the top), SCID tumors (n=5, labeled with a red bar at the top) and WT tumors (n=5, labeled with a blue bar at the top), measured by the NanoString nCounter system and analyzed using the NanoString nSolver tool. Each column represents one sample and each row represents a gene. **(F)** t-SNE plot of immune cells from WT (blue) and SCID (red) tumors. Live/CD45^+^ cells from WT and SCID tumors were used to generate a t-SNE map clustering cells based on the expression of their surface markers, as described in the Methods section (n=5 per group). **(G)** Cells in the t-SNE plot were sub-clustered using PhenoGraph algorithm. CD8^+^ T cell clusters in pink (Clusters A and B) and CD4^+^ T cell clusters in purple (Clusters C and D) (n=5 per group). **(H)** T cell clusters were further charachterized using ClusterExplorer analysis, representing the median fluorescence intensity of the selected markers in a heatmap format. One asterisk (*) indicates p value < 0.05, 2 asterisks (**) indicates p value < 0.01, 3 asterisks (***) indicates p value < 0.005, and 4 asterisks (****) indicates p value < 0.001.

To validate that the reduced mT3-2D tumor growth rate in WT mice is T cell-dependent, we depleted CD4^+^ and CD8^+^ T cells from immunocompetent mice. We observed that mT3-2D tumors in T cell depleted mice grew similarly to those in SCID mice (**Figure 1B** **and Supplementary Figure 3)**. Because missense mutations may generate T cell neoepitopes that can drive anti-tumor T cell responses, we performed whole exome sequencing (WES) on the mT3-2D cell line, 5 WT tumors and 5 SCID tumors. In total, we identified 1248 peptides with predicted binding affinities >500 nM. After filtering, 125 of these mutations were predicted to be putative cancer neoantigens **(Figure 1C)** (**Supplementary Table 1)**. Among the fourteen potential neoepitopes that were shared among the three groups, 13 belong to unique genes and 7 were validated by Sanger sequencing **(Supplementary Table 2)**. These observations suggest that the antitumor T cell response in WT mice was likely caused by the response to tumor neoantigens rather than by non-specific immune reactions.

To evaluate intratumoral T cell infiltration, we used flow cytometry and immunohistochemistry (IHC) analyses. We determined that there is a modest infiltration of T cells (**Figure 1D** **and Supplementary Figure 4A)**. To comprehensively assess the immune milieu of mT3-2D WT and SCID tumors, we employed NanoString nCounter immune profiling and multiparametric flow cytometry analysis. The NanoString heatmap analysis showed a clear separation of the WT and SCID samples and indicated greater upregulation of immune-related genes in mT3-2D WT tumors than in SCID tumors **(Figure 1E)**. In addition, NanoString nCounter Cell Type Score analysis showed that “T cell” and “cytotoxic cell” scores were significantly higher in the WT tumors **(Supplementary Figure 4B-C)**. In line with that observation, the analysis further indicated that interferon gamma (*Ifng*) expression was induced at significantly higher levels in the WT group than in the SCID group **(Supplementary Figure 4D)**. This was confirmed using RT-qPCR **(Supplementary Figure 4E)**.

Consistent with the NanoString nCounter analysis, a t-distributed stochastic neighbor embedding (t-SNE) analysis of multiparametric flow cytometry data indicated that cells from WT and SCID tumors were well-separated, suggesting that WT and SCID intratumoral immune profiles are notably different **(Figure 1F)**. In addition to t-SNE analysis, PhenoGraph analysis showed that intratumoral T cells were heterogenous and clustered into two CD8^+^ T cell subsets (Clusters A and B) and 2 CD4^+^ T cell subsets (Clusters C and D) **(Figure 1G)**. ClusterExplorer analysis was used to generate a heatmap analysis to determine the distinct phenotypes of the CD8^+^ and CD4^+^ T cell subsets **(Figure 1H)**, and thus are likely to possess different effector functions.

These findings suggest that mT3-2D WT and SCID tumors exhibit distinct immune profiles. Although T cell immunity is clearly induced in immunocompetent mice, T cell-based anti-tumor responses incompletely regulate mT3-2D tumor growth.

### Immune Selection Pressure Stimulates Malignant Epithelial Pancreatic Cells to Induce the Expression of Genes Associated with Myeloid Cells

We hypothesized that T cell immunity failed to completely control tumor growth because malignant pancreatic epithelial cells deploy immunosuppressive mechanisms. To investigate how these cells specifically respond to immune selection pressure, green fluorescent protein (GFP)-labeled mT3-2D cells were grown in C57BL/6J WT and SCID mice. RNA sequencing (RNA-seq) was used to examine the gene expression profiles of the mT3-2D-GFP cell line and FACS-sorted mT3-2D-GFP cancer cells isolated from the WT and SCID tumors. 52 genes were differentially expressed between the WT and SCID malignant epithelial pancreatic cells **(Figure 2A)**. A heatmap of the 52 differentially expressed genes is shown in **Figure 2B** and differentially expressed genes are listed in **Supplementary Table 3**. Among the genes that were upregulated in the FACS-sorted WT cancer cells compared to SCID cancer cells were chemokine (C-C motif) ligand 9 (*Ccl9*), arginase (*Arg1*) and cluster of differentiation 68 (*Cd68*). The selective upregulation of *Cl9*, *Arg1* and *Cd68* in the WT malignant cells was validated by RT-qPCR **(Figure 2C)**. Because these genes are typically expressed by myeloid cells, using RT-qPCR and flow cytometry, we determined that the FACS-sorted tumor cells do not express CD45 **(Supplementary Figure 5)**. This observation suggests that there was no apparent stromal contamination and no immune cell-cancer cell fusion. Also, we validated that the mT3-2D cell line as well as three additional murine KPC cell lines (mT4-2D, mT5-2D and KP1) possess basal expression of CCL9, ARG1 and CD68 at the mRNA and protein levels **(Supplementary Figure 6)**. Because the expression of *Cd68* was very low in FACS-sorted malignant cells, only *Ccl9* and *Arg1* were investigated in subsequent experiments. Together, these results suggest that malignant epithelial pancreatic cells express some genes associated with myeloid cells in response to immune selection pressure.

**Figure 2.**
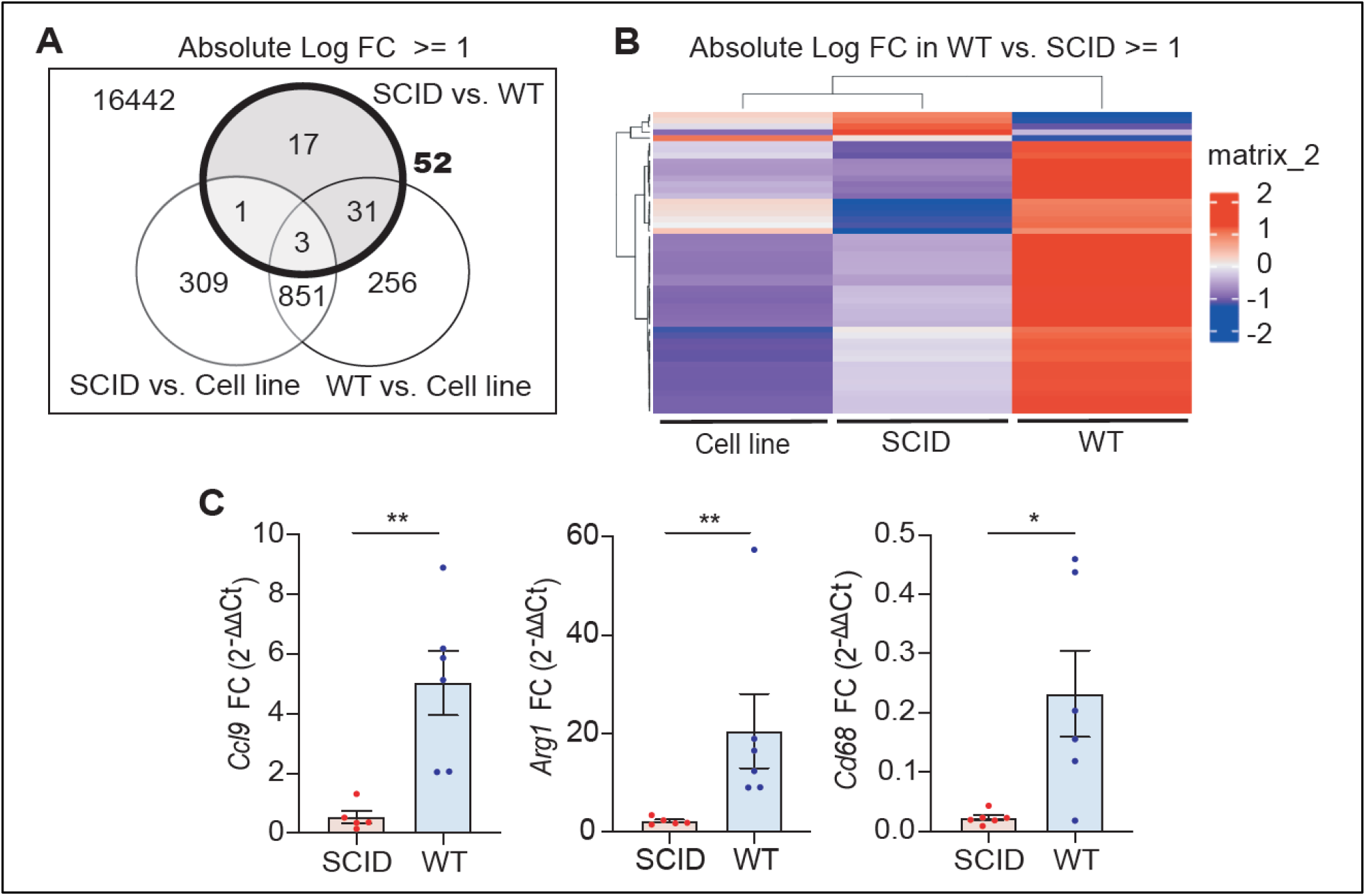
WT malignant epithelial pancreatic cells express higher levels of genes associated with myeloid cells compared to SCID malignant cells. **(A)** Number of differentially expressed genes between the mT3-2D-GFP cell line and tumor-derived FACS-sorted mT3-2D-GFP cells isolated from WT and SCID tumors identified by RNA-seq. Differential gene expression analysis was conducted for genes possessing an log2 fold change (FC) of at least 1. **(B)** Heatmap of the 52 differentially expressed genes between tumor-derived FACS-sorted mT3-2D-GFP cells isolated from WT and SCID tumors. Differential gene expression analysis was conducted for genes possessing an absolute log2 FC (fold change) of at least 1. Hierarchical clustering based on samples (columns) and genes (rows). **(C)** Quantitative real-time PCR analysis of *Ccl9*, *Arg1* and *Cd68* expression in FACS-sorted mT3-2D-GFP WT cancer cells (n=6) and FACS-sorted mT3-2D-GFP SCID cancer cells (n=5-6). Data are represented as mean FC (±SEM) and Mann-Whitney test was used. One asterisk (*) indicates p value < 0.05, and two asterisks (**) indicate p value < 0.01.

### Immune Selection Pressure is Associated with the Induction of Intratumoral PD-L1-Expressing Myeloid Cells

To determine the overall impact of immune selection pressure on mT3-2D tumors, including cancer cells and the surrounding stroma, we employed unbiased global proteomics and multiparametric flow cytometry analyses. Global proteomics profiling of WT and SCID whole tumors indicated that S100a8 and S100a9 proteins, which are typically associated with myeloid cells, were higher in the WT tumors than in the SCID tumors **(Figure 3A)**. We validated this observation using IHC **(Supplementary Figure 7A-B)**.

**Figure 3.**
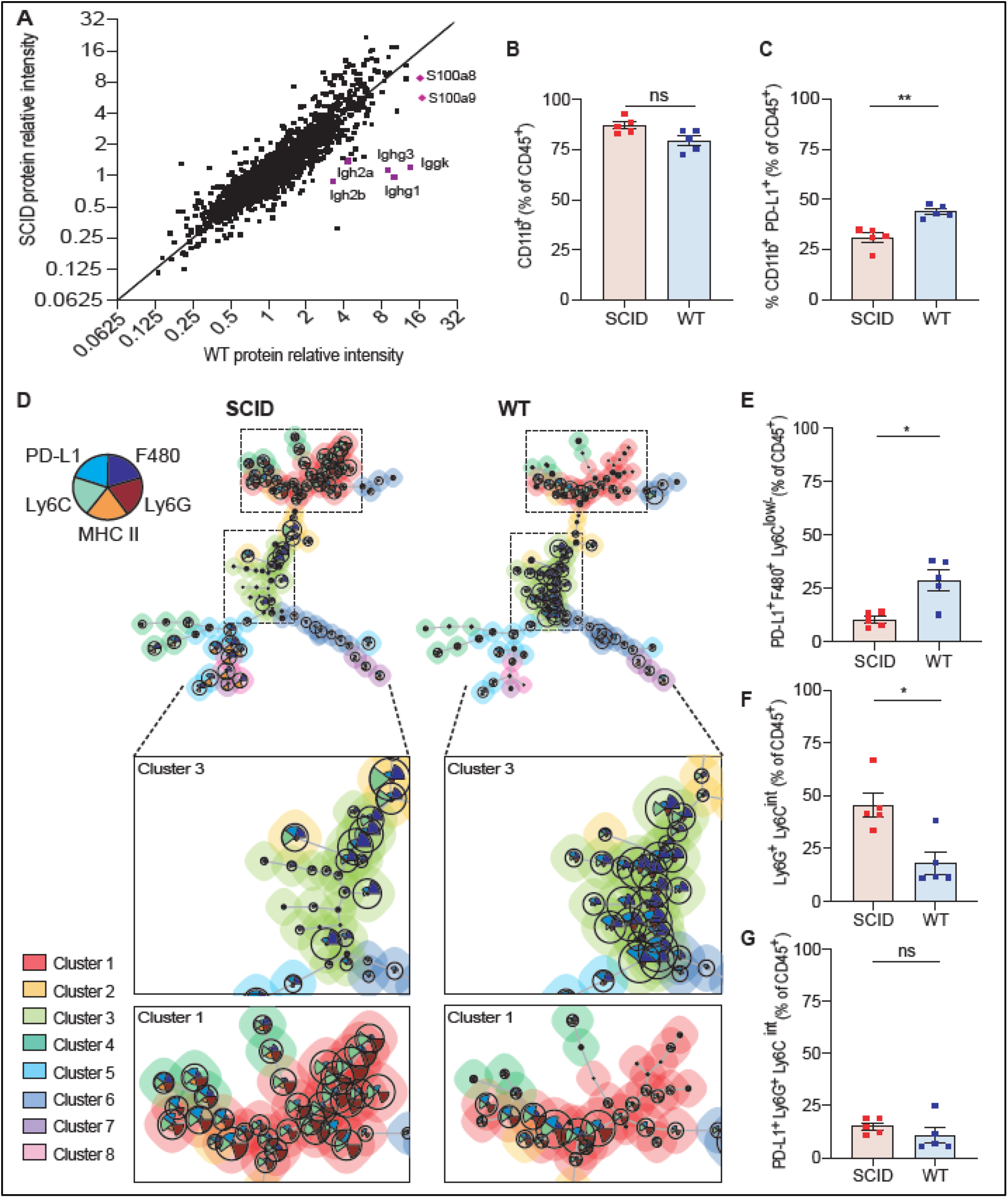
WT and SCID tumors have distinct myeloid cell profiles. **(A)** A scatter plot of paired relative protein intensities of WT and SCID tumors. Each dot represents a protein, and the identity line that passes through the origin is shown. **(B-C)** Percentage of CD11b^+^ cells in WT and SCID tumors detected by flow cytometry and represented as frequency of CD45^+^ cells using FlowJo (n=5 per group). Samples were gated as shown in **Supplementary Figure 7D-E**. A Mann-Whitney test was used for the analysis. **(D)** Multidimensional single-cell analysis was performed by FlowSOM analysis. Thirty thousand events of CD45^+^ cells from WT and SCID tumors were concatenated and then analyzed using FlowSOM (FlowJo), which identified 8 clusters (colored). Clustering was done based on the expression of Ly6G, Ly6C, PD-L1, F480 and MHC II. (**E-G**) Percentage of myeloid cell populations in WT and SCID tumors detected by standard gateing of flow cytometry data and represented as frequency of CD45^+^ cells using FlowJo (n=5 per group). Samples were gated as shown in **(Supplementary Figure 7D-E).** Mann-Whitney test was used for the analysis. ns=not significant, one asterisk (*) indicates p value < 0.05 and 2 asterisks (**) indicate p value < 0.01.

Flow cytometry analysis of the immune infiltrates determined that CD11b^+^ myeloid cells accounted for approximately 80% of total leukocytes in WT and SCID tumors, and there was no significant difference in the percentage of CD11b^+^ cells between the two groups (**Figure 3B** **and Supplementary Figure 7C)**. However, the frequency of CD11b^+^ cells expressing PD-L1 was significantly higher in the WT tumors **(Figure 3C)**. To better characterize intratumoral myeloid cell populations, we performed an unsupervised multidimensional single-cell analysis on CD45^+^ cells using the FlowSOM clustering approach. Based on the expression of Ly6C, Ly6G, F480, PD-L1 and MHC II, eight clusters were identified. Among these eight clusters, most cells segregated into 2 clusters. The cells segregated in Cluster 3 were F480^+^ PD-L1^+^ MHC II^low^, suggesting that they are PD-L1-expressing tumor-associated macrophages (TAMs). This cluster was predominantly noted in the WT group. The cells segregated in Cluster 1 were characterized by Ly6G^high^ and intermediate Ly6C expression (Ly6C^int^). This cluster was predominant in the SCID group. Both populations are magnified for visualization in **Figure 3D**. Interestingly, Cluster 1 contained heterogenous populations, with nodes containing PD-L1^+^ cells that were similar in WT and SCID tumors.

A standard flow cytometry gating strategy was used to confirm the FlowSOM observations **(Supplementary Figure 7E)**. Consistent with the FlowSOM analysis, the percentage of CD11b^+^ F480^+^ PDL1^+^ Ly6G^−^ Ly6C^low/−^ MHC II^−^ was higher in the WT tumors **(Figure 3E)**. The percentage of CD11b^+^ Ly6G^+^ Ly6C^int^ cells was higher in the SCID tumors **(Figure 3F)**, and there was no significant difference in the frequency of CD11b^+^ Ly6G^+^ Ly6C^int^ cells expressing PD-L1 between both groups **(Figure 3G)**.

The expression of CCL9 in cancer cells is known to be associated with immune evasion via the recruitment of C-C motif chemokine receptor 1 (CCR1)-expressing immunosuppressive myeloid cells into the tumors (10–14). Hence, we evaluated the expression of CCR1 in intratumoral CD11b^+^ cells using flow cytometry. We determined that the average of CD11b^+^ cells expressing CCR1 in WT tumors was 44.5% **(Supplementary Figure 8A-B)**. Also, to evaluate the expression of S100a8 and S100a9 in myeloid cell populations, we FACS-sorted CD11b^+^ Ly6G^−^ Ly6C^low/−^ and CD11b^+^ Ly6G^+^ Ly6C^int^ splenocytes, and then performed RT-qPCR. We determined that both myeloid cell populations expressed high levels of S100a8 and S100a9 **(Supplementary Figure 8C)**.

Collectively, these observations suggest that immune selection pressure altered intratumoral myeloid cell profiles. These alterations included induction of intratumoral PD-L1-expressing CD11b^+^ myeloid cells, potentially via CCL9/CCR1 signaling.

### Induction of STAT1 Expression in Immunocompetent Whole Tumors and Malignant Epithelial Pancreatic Cells

To identify the molecular determinants of immune evasion in mT3-2D tumors, differentially regulated genes and pathways between immunocompetent and immunodeficient tumors were examined. NanoString nCounter differential expression analysis identified nine genes that were upregulated in the WT whole tumors compared to the SCID whole tumors **(Figure 4A)**. The common upstream regulator of all of these genes, except chitinase-like protein 3 (*Chil3*), an M2 macrophage marker, is *Stat1* **(Figure 4B)**. The analysis of *Stat1* expression by NanoString nCounter analysis **(Figure 4C)**was validated by Western blot **(Figure 4D)**, and a similar pattern was shown by mass spectrometry **(Figure 4E)**.

**Figure 4.**
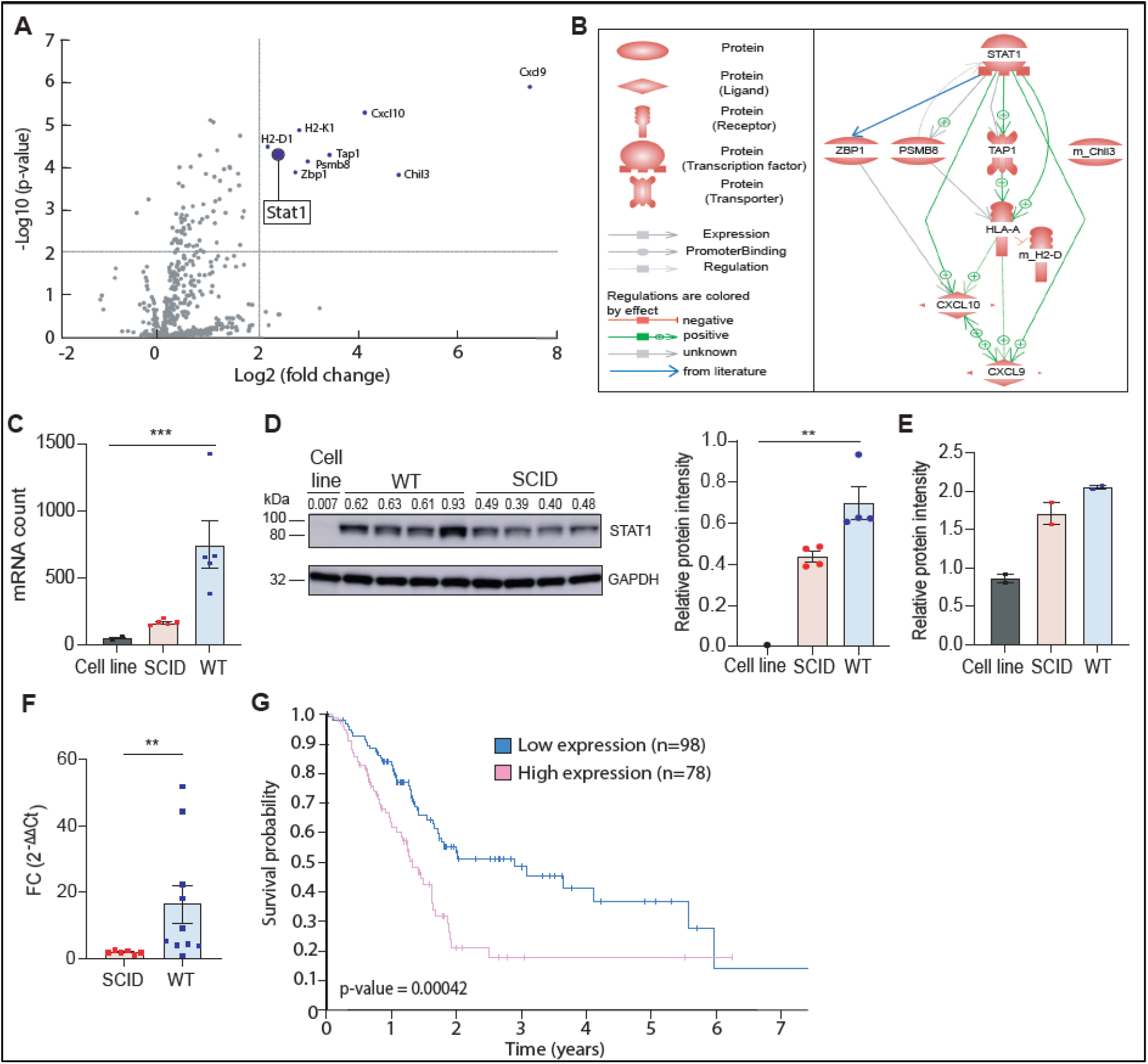
STAT1 expression is selectively more highly induced in immunocompetent tumors than in immunodeficient tumors. **(A)** Volcano plot of NanoString nCounter differential gene expression of all pairwise comparisons. The plot displays the relationship between fold-change and p-value between the WT and SCID groups using a scatterplot view. Each dot represents the average expression of a gene in the WT samples in reference to the SCID samples. **(B)** Direct interaction analysis by Pathway Studio for eight out of the nine differentially expressed genes identified in **(A)**. Interaction searching parameters are promoter binding, expression and regulation. The Pathway Studio figure was adapted by the authors to include a blue arrow that shows the link between STAT1 and ZBP1 that was reported in the literature but was not identified by Pathway Studio (50). **(C)** *STAT1* RNA counts as determined using the NanoString nSolver tool (n=2 technical replicates for the cell line, n=5 biological replicates for the tumor samples). Data represented as mean (±SEM) and a Kruskal-Wallis test was used. **(D)** Western blot of STAT1 and GAPDH in the mT3-2D cell line, 4 WT tumor samples and 4 SCID tumor samples. Densitometry values of expression relative to GAPDH are indicated above the western blot and shown in the bar plot to the right. Data represented as mean (±SEM) and a Kruskal-Wallis test was used. **(E)** Relative STAT1 protein intensity detected by mass spectrometry. **(F)** Quantitative real-time PCR analysis of *STAT1* expression in FACS-sorted mT3-2D-GFP WT cancer cells (n=11) and FACS-sorted mT3-2D-GFP SCID cancer cells (n=6). Data represented as mean fold change (FC) (±SEM), and a Mann-Whitney test was used. **(G)** Kaplan-Meier plot showing *STAT1* pancreatic cancer survival using the TCGA data from the Human Protein Atlas. The x-axis shows time for survival (years) and the y-axis shows the probability of survival, where 1.0 corresponds to 100 percent. The cutoff is 33.2, the 5-year survival for patients with higher expression than the expression cutoff (high expression) is 18% and the 5-year survival for patients with lower expression than the expression cutoff (low expression) is 37%. One asterisk (*) indicates p-value < 0.05, 2 asterisks (**) indicate p-value < 0.01 and 3 asterisks (***) indicates p value < 0.005.

In addition to whole tumor analysis, RNA-seq Hallmark pathway analysis of the mT3-2D-GFP cell line and FACS-sorted mT3-2D-GFP cancer cells isolated from WT and SCID tumors was used to identify pathways that are modified specifically in the malignant cells. Interferon-gamma-response and interferon-alpha-response were the most significantly enriched pathways in the FACS-sorted mT3-2D-GFP cancer cells isolated from tumors grown in WT mice as compared to tumors grown in SCID mice (FDR adjusted p-values of 1.72×10^−11^ and 1.29×10^−06^, respectively) **(Supplementary Table 4)**. Malignant epithelial cell analysis also determined that *Stat1* was selectively induced in the WT malignant cells (**Supplementary Table 3)**, and this finding was validated by RT-qPCR **(Figure 4F).** Using the Cancer Genome Atlas (TCGA) data from the Human Protein Atlas, it was found that high *STAT1* expression is significantly associated with shorter overall survival for all pancreatic cancer patients **(Figure 4G)**, suggesting a potential role for STAT1 in human pancreatic cancer immune evasion.

Notably, differences in the induction of other *Stat* genes *(Stat2, Stat3, Stat4, Stat5b,* and *Stat6)* were less prominent between WT and SCID whole tumors compared to *Stat1* by NanoString nCounter analysis **(Supplementary Figure 9A-E)**. Similarly, using mass spectrometry, none of the other STAT proteins was selectively induced in the WT whole tumors compared to the SCID whole tumors **(Supplementary Figure 9F)**. Also, Human Protein Atlas survival analysis indicated that while *STAT2, STAT3, STAT5A* and *STAT6* were not significant prognostic factors, high *STAT4* and *STAT5B* expression was associated with longer overall survival in pancreatic cancer **(Supplementary Figure 10)**. Together, these results suggest that STAT1 is selectively induced both in WT malignant cells and in WT tumor microenvironments compared to SCID samples, and that STAT1 may play a role in immune evasion.

### Ruxolitinib downregulates the expression of CCL9 and ARG1 and increases the anti-tumor efficacy of anti-PD1 in immunocompetent mice

It has been reported that STAT1 signaling regulates TAM-mediated T cell inhibition by regulating arginase activity and nitric oxide production (15). Also, CCL9 expression is strongly induced by the interferon consensus sequence-binding protein (ICSBP) (16). To investigate the effect of JAK/STAT signaling on the expression of *Ccl9* and *Arg1*, we treated mT3-2D cells *in vitro* with ruxolitinib for up to 12 hours. Using RT-qPCR, we determined that the expression of *Ccl9* and *Arg1* were downregulated upon ruxolitinib treatment **(Supplementary Figure 11A-B)**. Also, the effect of ruxolitinib on *Ccl9* and *Arg1* was not unique to mT3-2D cells, as the effect was reproduced in multiple murine KPC cell lines **(Supplementary Figure 11C-D)**.

To determine if the ruxolitinib-mediated inhibition of CCL9 and ARG1 was maintained *in vivo* and if ruxolitinib had anti-tumor effects, we treated mT3-2D immunocompetent mice with either PBS, ruxolitinib, anti-PD1 or the combination of anti-PD1 and ruxolitinib. Although ruxolitinib monotherapy did not result in a significant tumor growth inhibition, ruxolitinib improved the anti-tumor efficacy of anti-PD1 treatment. In fact, although anti-PD1-treated mice had an initial reduction in tumor size, the majority of the tumors eventually progressed. However, all tumors in the combination group were completely rejected **(Figure 5A**, **Supplementary Figure 12)**. Using Western blot, CCL9 protein level was reduced in tumors from the ruxolitinib-treated mice compared to the control group, which was consistent with the expression of S100a9 (S100a8 was not detectable by Western blot in both groups). A similar but less consistent pattern of inhibition was also seen in ARG1 protein expression **(Figure 5B)**. Of note, ruxolitinib did not affect the cellular proliferation rate of mT3-2D cells *in vitro* even at high concentrations **(Supplementary Figure 13A)**, suggesting that the ruxolitinib effect is host immunity-dependent. Knowing that JAK/STAT signaling is essential for immune cell differentiation and function (17), the impact of ruxolitinib *in vivo* on T cell number and activity was evaluated using flow cytometry. There were no significant differences in the frequency of intratumoral or spleen CD4^+^ and CD8^+^ T cells **(Supplementary Figure 13B)**. Also, there were no notable differences in the percentages of CD8^+^ T cells expressing IFNγ between the two groups (7.3% in 3 combined PBS-treated tumors and 8.8% in 3 combined ruxolitinib-treated tumors), and in CD4^+^ T cells expressing IFNγ (5.7% in 3 combined PBS-treated tumors and 6.6% in 3 combined ruxolitinib-treated tumors).

**Figure 5.**
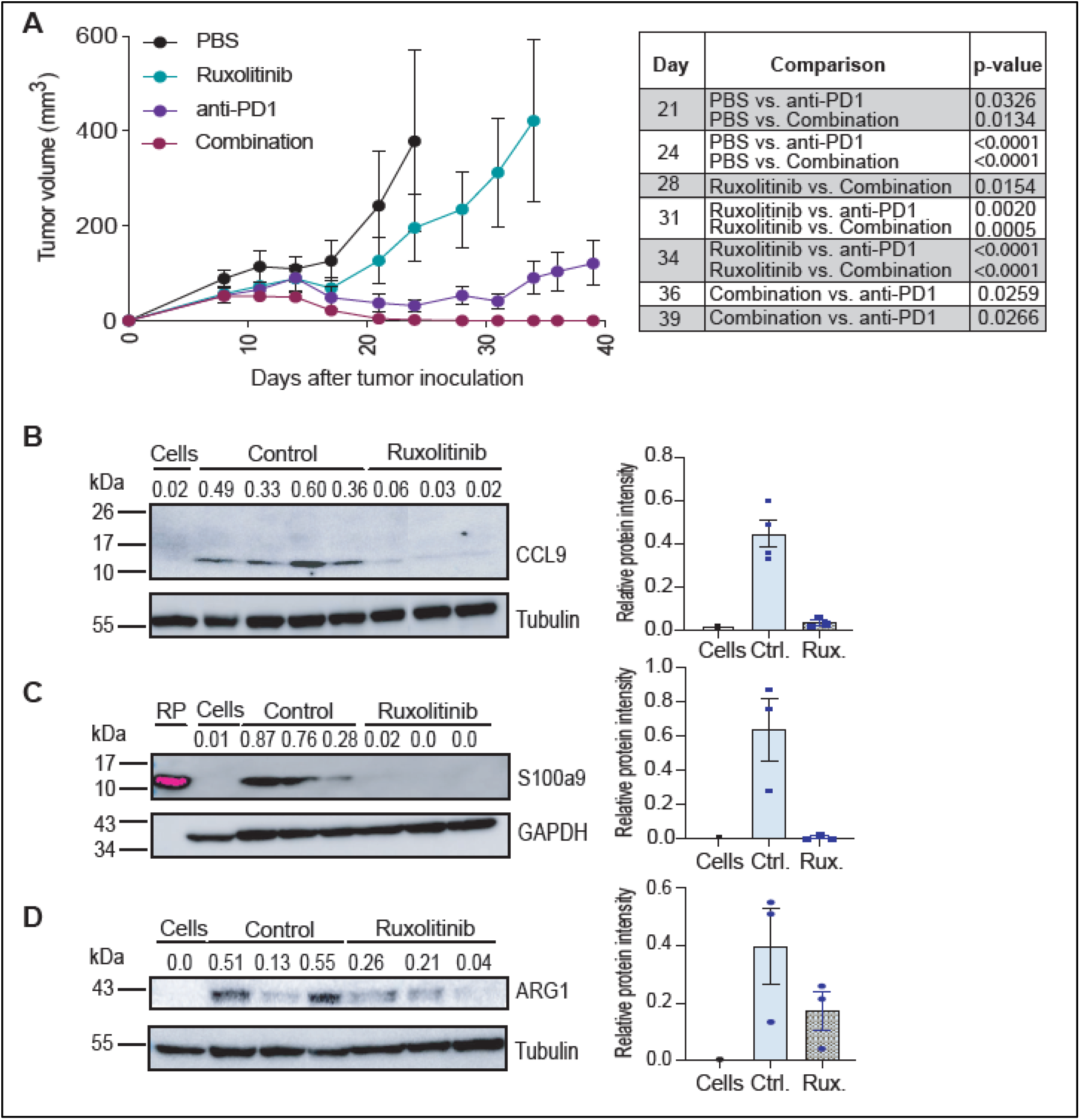
Ruxolitinib downregulates CCL9 and ARG1 expression *in vivo* and improves the anti-tumor efficacy of anti-PD1 treatment. **(A)** Tumor growth curve depicting the average (±SEM) volume of mT3-2D tumors grown in C57BL/6J immunocompetent mice treated with PBS (n=5), ruxolitinib (n=5), anti-PD1 (n=7) and combination of anti-PD1 and ruxolitinib (n=5). A two-way ANOVA followed by Tukey multiple comparison test was used for up to day 34, and two-tailed t-test was used for days 36 and 39. **(B-D)** Western blot of CCL9, tubulin, S100a9, ARG1 and GAPDH in the indicated samples. Densitometry values of expression relative to either tubulin or GAPDH indicated above the western blot and shown in the bar plot to the right. (RP=S100a9 recombinant protein, R&D systems, #8916-S8-050). Data represented as mean (±SEM). (Rux.= ruxolitinib, Ctrl.= control)

## Discussion

The goal of this study was to determine the extent to which immune selection pressure shapes the immunosuppressive tumor microenvironment in a murine model of pancreatic ductal adenocarcinoma. Here, we have demonstrated that, despite the clear induction of T cell responses in syngeneic mT3-2D pancreatic ductal adenocarcinoma-bearing immunocompetent WT mice, T cell immunity incompletely regulates tumor growth. Immune selection pressure induces a myeloid mimicry phenomenon by malignant epithelial pancreatic cells, stimulating intratumoral infiltration of PD-L1-expressing myeloid cells. These tumor cell-based responses to immune selection pressure promote immune escape in this model system, which can be reversed by inhibiting JAK/STAT signaling **(Figure 6)**.

**Figure 6.**
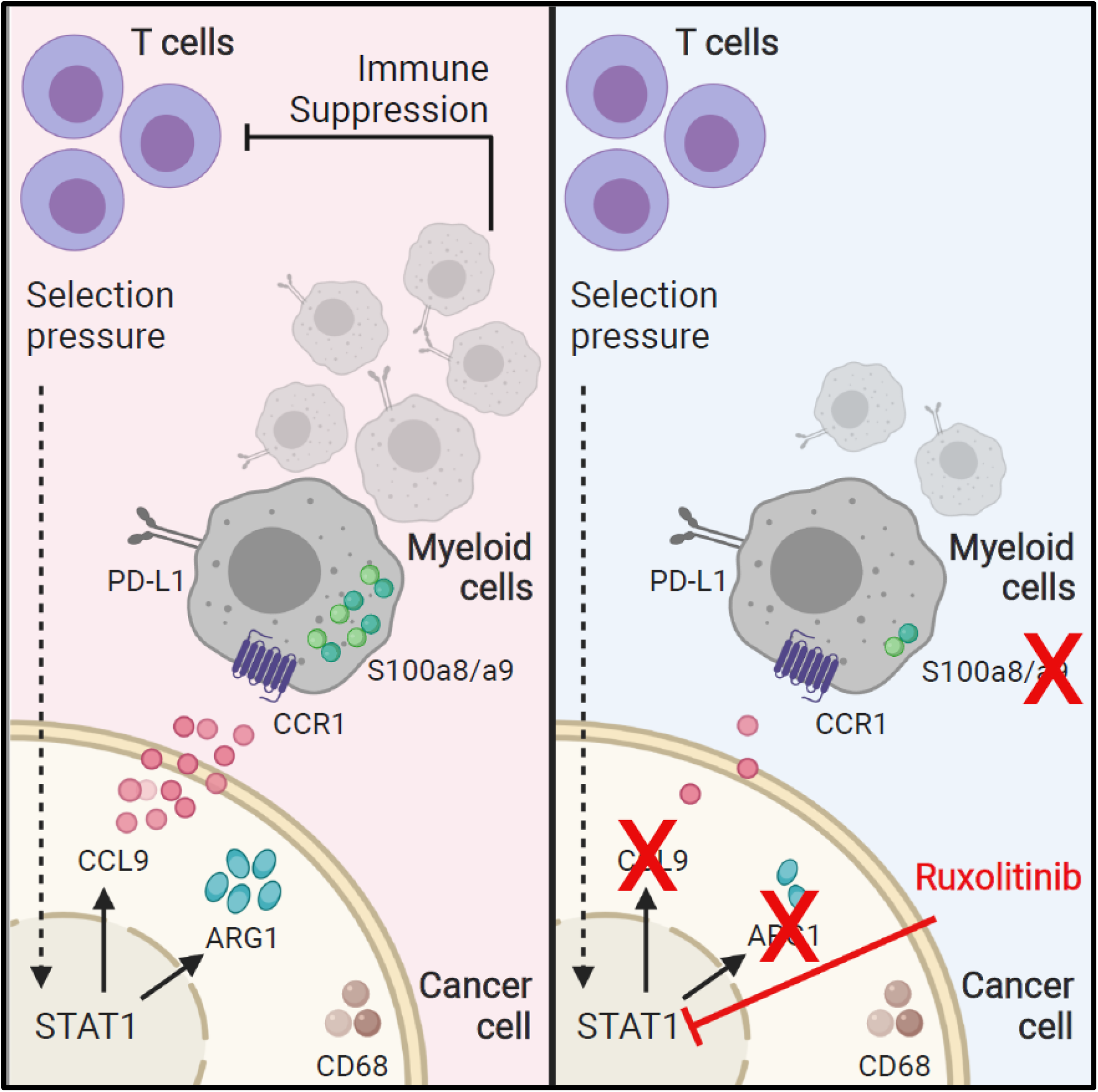
Immune selection pressure induces pancreatic cancer immune suppression. Graphical representation of the linkages between immunogenicity and tumor-derived immunosuppression. Targeting JAK/STAT signaling, using ruxolitinib, reduces the expression of CCL9, S100a9 and ARG1.

T cells within human pancreatic tumors are generally infrequent and incapable of eliciting anti-tumor immunity (3,9). Thus, T cell immunity has been considered irrelevant to pancreatic cancer, and it has been argued that the pancreas might have organ site-specific resistance to immunotherapy. This hypothesis was supported by the observation that the vast majority of pancreatic cancer immunotherapy clinical trials have failed (9). However, recent studies show that tumors of long-term pancreatic cancer survivors exhibit potent cytotoxic T cell responses (7). Also, immune checkpoint inhibitors, which promote T cell responses, have demonstrated efficacy in pancreatic cancer patients with MSI-high tumors (8). Hence, T cell immunity is likely to play a critical role in pancreatic cancer biology, but further resolution of the molecular determinants of immune evasion in pancreatic malignancy is needed. The goal of this study was to address the extent to which immune evasion in pancreatic cancer is a tumor-intrinsic property and if it is induced in response to immune selection pressure. If the latter is true, identifying immunosuppressive mechanisms that cancer cells employ in order to survive in the face of T cell recognition and attack may facilitate the development of novel immunotherapies and improve responses to currently available treatment strategies.

To address this goal, we investigated the effects of immune selection pressure on pancreatic cancer using the mT3-2D subcutaneous murine model, which is syngeneic to the C57BL/6 strain. Two types of analysis were performed: (i) malignant epithelial cell analysis, and (ii) whole tumor analysis. The former determines how malignant cells specifically respond to immune selection pressure, whereas the latter determines the effect of immune selection pressure on the tumor microenvironment, including cancer cells and non-cancer stromal elements. Combining these two approaches provided a comprehensive understanding of the complex interactions between cancer cells and the immune system.

Using another KPC murine model, Evans et al. reported that depleting T cells during the development of spontaneous tumors did not affect tumor natural history. Using neoantigen prediction computational algorithms, these tumors were determined to lack immunogenic neoepitopes. However, the introduction of immunogenic antigens into these KPC tumors stimulated T cell anti-tumor responses (18). In contrast, in the mT3-2D subcutaneous mouse model used here, we identified 125 putative cancer neoantigens within 11 samples (mT3-2D cell line, 5 WT tumors and 5 SCID tumors), with 14 potential neoepitopes predicted to be shared between the three groups, half of which were validated by Sanger sequencing. The slow tumor growth rate in immunocompetent mice suggests that at least some of these mutations stimulated T cell immunity. These findings are consistent with those reported by Boj et al. in the mT3-2D model (19) and by Balachandran et al. and Bailey et al. in human pancreatic adenocarcinomas (7,20). Hence, despite the expression of some potentially immunogenic antigens in the malignant cells, pancreatic tumors employ mechanisms to evade cytotoxic T cell immunity.

The myeloid mimicry phenomenon in cancer cells has been reported previously in several cancer entities, including breast cancer, colorectal cancer and pancreatic cancer, and it is associated with advanced disease and worse outcomes (21–23). Some studies determined that the expression of myeloid markers by cancer cells was a result of cancer cell-immune cell fusion, and thus created hybrids that express characteristics of both cellular origins (24–26). In our study, we determined that FACS-sorted malignant epithelial cells and murine KPC cell lines express chemokine (C-C motif) ligand 9 (CCL9), arginase 1 (ARG1), and cluster of differentiation 68 (CD68), but do not express CD45, the transmembrane protein expressed by all hematopoietic cells, suggesting that the myeloid mimicry phenomenon we reported here is not a consequence of cell fusion. Our data suggest that immune selection pressure plays an important role in regulating this phenomenon.

Whole tumor analysis using global proteomics profiling and multiparametric flow cytometry analysis revealed selective increases of the myeloid cell-associated proteins S100a8 and S100a9, and increased infiltration of PD-L1^+^ myeloid cells in the WT as compared to SCID tumors, respectively, potentially via CCL9/CCR1 signaling. CCL9 induction by cancer cells has been shown to be associated with the recruitment of CCR1-expressing immunosuppressive myeloid cells, and that targeting CCL9/CCR1 interaction limited tumor growth in mice. However, the phenotype of these CCR1-expressing myeloid cell populations varies among cancer models (10–14). Together, our findings suggest that immune selection pressure, potentially through myeloid mimicry, was associated with the infiltration of PD-L1-expressing myeloid cells that contributes to establishing an immunosuppressive pancreatic tumor microenvironment.

Furthermore, whole tumor analysis consistently indicated upregulation of STAT1 protein in WT whole tumors compared to SCID whole tumors. However, other STAT proteins were less notably different between the two groups. Also, we found that STAT1 expression was upregulated selectively in epithelial cells isolated from WT tumors, suggesting a role for STAT1 as an orchestrator of this myeloid mimicry phenomenon. Interestingly, a recent study has identified an interferon (IFN)-stimulated gene (ISG) resistance signature (ISG.RS), predominantly expressed in human cancer cells, and associated with resistance to immune checkpoint blockade. *STAT1*, but not any other *STAT* genes, is a component of this ISG.RS signature (27). We found that JAK/STAT signaling regulates the expression of CCL9 and ARG1, which are reduced in mT3-2D cells following ruxolitinib exposure. Although *in vivo* ruxolitinib treatment targets malignant cells as well as the tumor stroma, the remarkable reduction of CCL9 in tumors from ruxolitinib-treated mice in comparison to control tumors suggests that the ruxolitinib effect targets the malignant cells. Muthalagu et al. recently described a role for mutant KRAS and MYC cooperation in suppressing STAT1 signaling in a murine model of pancreatic cancer (28). In contrast to our study, they did not examine the impact of immune selection on these molecular interactions. Moreover, STAT1 overexpression is an adverse prognostic factor in human pancreatic cancer. Together, these data suggest that potentially immunosuppressive CCL9 and ARG1 expression in cancer cells were elicited in response to immune selection pressure through JAK/STAT signaling.

We found that ruxolitinib treatment improves anti-PD1 therapeutic effects in immunocompetent mice bearing mT3-2D subcutaneous tumors. A similar observation was reported in a recent study by Lu et al. using the PANC02 orthotopic murine pancreatic cancer model (29). This study also reported that ruxolitinib induced cytotoxic T cell frequency and activity in the tumor microenvironment. Hence, our findings may be generally applicable.

Myeloid cell populations are highly heterogenous (30,31). While many myeloid cell populations can stimulate immune evasion via a wide range of immunosuppressive mechanisms, some myeloid cells have anti-tumor properties (32). However, PD-L1 expressing myeloid cells have been reported to promote immune escape (33,34). Thus, our observations suggest that immune selection pressure alters intratumoral myeloid cell profiles that likely contribute to immune evasion in pancreatic ductal adenocarcinoma.

Despite the induction of tumor-directed T cell immunity in immunocompetent mice bearing mT3-2D tumors, T cell immunity incompletely controlled tumor growth. We have shown that immune selection pressure, via JAK/STAT signaling, stimulated a myeloid mimicry phenomenon, and this was associated with the intratumoral infiltration of PD-L1-expressing myeloid cells that likely play an important role in immune evasion. These observations could be relevant to human pancreatic cancer and potentially other malignancies. Future studies will further elucidate the specific role of STAT1 signaling in orthotopic murine pancreatic cancer models and in genetically engineered mouse models that permit study of the influence of immune selection pressure on the evolution of tumor-derived immunosuppression.

## Methods

### Cell Lines

mT3-2D, mT4-2D and mT5-2D murine pancreatic cancer cell lines were gifts from David Tuveson, Cold Spring Harbor Laboratory, Laurel Hollow, NY (19). The KP1 murine pancreatic cancer cell line was a gift from David DeNardo, Washington University Medical School, St. Louis, MO (35). PANC02 murine pancreatic cancer cells were a gift from Jill Smith, Georgetown University Lombardi Comprehensive Cancer Center, Washington, DC. mT3-2D GFP cell line was a gift from Chunling Yi, Georgetown University Lombardi Comprehensive Cancer Center, Washington, DC. Briefly, pHAGE PGK-GFP-IRES-LUC-W (addgene, #46793) was transfected into 293T cells to generate the virus. The virus was infected into mT3-2D cells and GFP positive cells were FACS-sorted. All these cell lines are syngeneic in C57BL/6 mice. HEK293T cells, C3H/10T 1/2 Clone 8 and RAW 264.7 cell lines were obtained from the Georgetown Lombardi Comprehensive Cancer Center Tissue Culture Shared Resource (TCSR), Georgetown University, Washington, DC). All cell lines were grown in standard conditions and maintained in Dulbecco’s Modified Eagle Medium (DMEM) (Fisher Scientific, # SH30022LS) supplemented with heat-inactivated fetal bovine serum (HI-FBS) to a final concentration of 10% and 2 mM L-glutamine, except C3H/10T 1/2 Clone 8 was cultured in Eagle’s Basal medium (Thermofisher, #21010-046) supplemented with HI-FBS to a final concentration of 10% and 2 mM L-glutamine. All cell lines were tested and determined to be free of Mycoplasma and other rodent pathogens.

### Animal Xenograft Studies

1 × 10^5^ of mT3-2D and mT3-2D-GFP cells were injected subcutaneously in the right flank of C57BL/6J wild-type (WT) and B6.CB17-Prkdc^scid^/SzJ (SCID) mice. 2 × 10^6^ of PANC02 cells were subcutaneously injected in the right flank of WT and SCID mice. All mice used in the study were 6–8 weeks of age. WT and SCID mice were purchased from The Jackson Laboratory (Bar Harbor, ME). For ruxolitinib (LC laboratories, Woburn, MA) *in vivo* treatment, WT mice were treated by oral gavage at a dose of 50 mg/kg daily, starting when tumors reached about 50-100 mm^3^ and continued for 4 weeks. Anti-PD1 (clone: RMP1-14, BioXcell) was used at a dose of 200 μg/injection twice per week. Anti-PD1 treatment started when tumors reached about 50-100 mm^3^ and continued for 4 weeks. Tumors were measured twice-weekly using calipers. Volume was calculated as (length × width^2^)/2. When tumors reached 1 cm^3^ or when mice showed signs of pain or distress, mice were euthanized using CO2 inhalation, and the tumors were excised. After euthanizing the mice, tissue samples were collected for downstream analysis. All studies involving animals were reviewed and approved by the Georgetown University Institutional Animal Care and Use Committee (GU IACUC).

### *In vivo* T Cell Depletion

1 × 10^5^ of mT3-2D cells were injected subcutaneously in the right flank of C57BL/6J mice. The *in vivo* depletion of CD4^+^ and CD8^+^ T cells, was done as described previously (36).

### Preparation of Solid Tumor Tissue for Single-Cell Suspension

Tumors were chopped into small pieces that were then transferred into gentleMACS tubes (MACS Miltenyi Biotec), containing 10 ml of DMEM media and 1 mg/ml collagenase D (Sigma-Aldrich, COLLD-RO Roche, #11088866001). The tubes were placed on a gentleMACS Dissociator (MACS Miltenyi Biotec, #130-095-937) using the program 37_m_TDK2. After incubation, cells were filtered using 70 μm cell strainer and recovered by centrifugation.

### Preparation of Splenocytes for Single-Cell Suspension

Spleen tissue was gently ground between frosted glass microscope slides. Tissue was passed 10x through a 1000 μL pipette tip. Red cells were removed by incubating the splenocytes for 3 minutes with 3 ml eBioscience™ 1X RBC Lysis Buffer (Invitrogen, ThermoFisher, #00-4333-57). Cells were pelleted by centrifugation, and then recovered in 10 ml RPMI media with 10% HI-FBS, and filtered using a 70 μm cell strainer.

### NanoString nCounter Mouse PanCancer Immune Profiling Screen

RNA was extracted from 5 WT tumor samples, 5 SCID tumor samples and the mT3-2D cell line using the PureLink RNA Mini Kit (Ambion). RNA concentration and quality were assessed using the Agilent BioAnalyzer. 100 ng of total RNA from each sample was hybridized to the CodeSet nCounter PanCancer Immune Profiling Panel, Mouse, and processed on the NanoString Technologies nCounter Sprint Profiler according to NanoString protocols in the Georgetown University Genomics and Epigenomics Shared Resource (GESR). nSolver software was used for data normalization and analysis.

### Whole Exome Sequencing (WES) and Neoantigen Prediction

DNA was extracted from 5 WT mT3-2D subcutaneous tumors, 5 SCID mT3-2D subcutaneous tumors and the mT3-2D cell line using a DNeasy Blood and Tissue Kit (Qiagen, #69504). Indexed, paired-end whole exome libraries were constructed from 1.0 μg gDNA using the Agilent SureSelect XT HS Mouse All Exon Library Kit. Libraries were sequenced on an Illumina HiSeq 4000 using PE 150 bp (300 cycle) chemistry to a mean depth of 200x per sample. For Sanger sequencing, Genewize (www.genewiz.com) was used to design primers that target the neoepitope regions. Then, each targeted region was amplified with PCR and sequenced with Sanger sequencing.

Reads were aligned to the mm10 reference genome using the Burrows–Wheeler Alignment tool version 0.7.17 (37). Genome Analysis Toolkit (GATK) best practices were used for sorting, duplicate marking, and variant calling with the HaplotypeCaller (38). Variants were annotated using Ensemble-VEP version 90.9 (39) and all possible 9-mer peptides that include a missense variant were identified using pVACtools version 1.0.8 (40). MHC-I binding affinities (H-2-Db and H-2-Kb) were then predicted for each wild-type and corresponding mutant peptide with NetMHC 4.0 (41). In addition, the following filtering steps were performed. Mutant peptides with predicted binding affinity > 500 nM were removed. Peptides from olfactory genes were excluded because they are highly prone to somatic mutations, are not highly expressed, and yield false positives as cancer-associated genes (42). Mutations that corresponded to known polymorphic genes were also excluded. Peptides with low predicted agretopic indices (43), i.e., ratio of mutant to wild-type binding affinity < 4, were removed.

### RNA-seq

Six mT3-2D-GFP WT and 5 SCID tumors were processed into single cell suspensions as described above. These cells were stained with Helix NP Blue (Biolegend, #425305) and live GFP positive cells were isolated using a BD FACSAria IIu fluorescence-activated cell sorting (FACS) cytometer. FACS-sorted cells were then snap frozen in liquid nitrogen. RNA was extracted from WT GFP positive FACS-sorted cells, SCID GFP positive FACS-sorted cells and the mT3-2D-GFP cell line using the RNeasy Micro Kit (Qiagen). RNA concentration and quality were assessed using an Agilent BioAnalyzer. Equal amounts of RNA from each corresponding group were combined as one sample (1 WT sample, 1 SCID sample and 1 cell line). Indexed, paired-end sequencing libraries were prepared from 1.0 μg total RNA using the Illumina TruSeq Stranded mRNA Library Preparation Kit (polyA cDNA synthesis) in the Genomics and Epigenomics Shared Resource (GESR) in the Lombardi Comprehensive Cancer Center at Georgetown University Medical Center. Libraries were sequenced on an Illumina NextSeq550 using 75 bp (150 cycle) paired-end chemistry to a mean depth of 40M reads per sample. Raw reads were aligned to the hg19 reference genome with BWA (37,44), and then quantified into read counts with the annotation model (Partek E/M). All subsequent RNA-seq analyses were performed in R version 3.5.1. Read counts were variance stabilized with the regularized log (rlog) function in the R/Bioconductor package DESeq2 version 1.20.0 (45). Log fold changes were computed on the rlog-transformed expression data and genes with absolute log fold changes greater than or equal to one were retained as significant. One-sided gene set statistics were performed for log fold changes on Hallmark version 6.2 pathways from MSigDB (46) and KEGG pathways (47) from the R/Bioconductor package KEGG.db version 3.2.3 and sets with FDR adjusted p-values below 0.05 were called statistically significant.

### Sample Preparation for Global Proteomics Profiling

Solid tumor tissue was ground with a mortar and pestle in the presence of liquid nitrogen to create a powdered tissue. The powdered tissue or cell pellets were resuspended in lysis buffer containing 50 mM Tris HCl pH 7.5, 100 mM NaCl, 1% Triton X-100, 10 mmol/L EDTA, 1 mmol/L phenylmethylsulfonyl fluoride (PMSF), 1× Protease Inhibitor Cocktail (Roche, Cat. No. 04693132001) and 1x Phosphatase Inhibitor Cocktail (Sigma, Cat. No. P5726). The suspension was vortexed vigorously for 5 minutes, sonicated and then maintained in agitation for 30 minutes at 4°C. The suspension was spun down at 12,000 x g for 15 minutes. The supernatant was collected, and samples were processed for NanoUPLC-MS/MS and analyzed as described previously (48).

### Western Blotting

Solid tumor tissue was ground with a mortar and pestle in the presence of liquid nitrogen to create a powdered tissue. Lysis buffer was added to the powdered tissue or cell pellets and maintained in agitation for 30 minutes at 4°C. The lysis buffer consists of RIPA Buffer (Thermo Scientific™, #89900), 10x Protease Inhibitor (Sigma, #S8820) and 1x Phosphatase Inhibitor Cocktail (Sigma, #P5726-1ML). Western blot was done as described previously (36). The antibodies are listed in **Supplementary Table 5**.

### RNA Isolation and Quantitative Reverse Transcription PCR (RT-qPCR)

RNA extraction and RT-qPCR were performed as described previously (36). Gene expression was normalized to HPRT or GAPDH and analyzed using the ΔCт or 2^−ΔΔCт^ method in triplicate. The primer sequences are listed in **Supplementary Table 6**. For lipopolysaccharide (LPS) treatment, RAW264.7 cells were treated with 1 μg/ml LPS (Sigma-Aldrich, # L3129) for 24 hours. For *in vitro* ruxolitinib treatment, cells were treated with 1 μg/ml of ruxolitinib (MCE, #HY-50856) for the indicated time periods.

### Immunohistochemistry (IHC)

For solid tumors, tissues were fixed in 10% formalin overnight at room temperature, and then stored in 70% ethanol until paraffin embedding. For HistoGel sample preparation, 1-5 million cells were fixed in 10% formalin overnight at 4°C. Cells were washed with PBS and then left in HistoGel for a few minutes, followed by the addition of 100% ethanol. Samples were sent to the Georgetown University Histopathology and Tissue Shared Resource for embedding, sectioning and staining. ImageJ (v1.48) and FIJI (v2.0.0-rc-69/1.52n) were used for the analysis. The antibodies are listed in **Supplementary Table 7**.

### Arginase Activity Assay

Arginase activity was measured colorimetrically using Abcam’s Arginase Activity Assay Kit (#ab180877). 5×10^6^ cells were lysed with 100 μl of the kit’s lysis buffer (2×10^6^ cells per well). Samples were processed and analyzed per the manufacturer’s instructions.

### Crystal Violet Staining

Crystal violet staining was done as described previously (36).

### Multiparametric Flow Cytometry Analysis of Intratumoral Immune Cell Infiltrates

Single cell suspensions were generated as outlined above. 2×10^6^ cells were prepared for staining as described previously (36). Then, cells were stained with a cocktail of surface mAbs: anti-mouse-CD45 (clone 30-F11, #564590), NK-1.1(clone PK136, #562864), B220 (clone RA3-6B2, # 563103), CD3e (clone 145-2C11, #564661), CD4 (clone RM4-5, #563151), CD8a (clone 53-6.7, #564920), PD1 (clone J43 #744549), PD-L1 (clone MIH5, #564715), CD25 (clone PC61, #565134), I-A/I-E (clone M5/114.15.2, #563415), CD11b (clone M1/70, #564443), CD11c (clone N418, #565452), all from BD Biosciences, and F480 (BM8, #123114), Gr-1 (clone RB6-8C5, #108428), Ly6G (Clone 1A8; #127645) and Ly6C (Clone HK1.4, #128030) from BioLegend. After 30 minutes of staining, cells were washed and samples were acquired with a FACS Symphony cytometer (BD Biosciences).

For the analysis, cells were manually gated on size and granularity. Dead cells and doublets were excluded, and CD45^+^ cells were selected **(Supplementary Figure 7d)**. Then, three thousand CD45^+^ live cells from each WT (n=5) and SCID mouse (n=5) were concatenated. A total of 30,000 CD45^+^ cells were analyzed using unsupervised t-SNE, FlowSOM, ClusterExplorer and PhenoGraph algorithms using FlowJo software.

For FlowSOM (Flow Self-Organizing Map, FlowJo) analysis, Ly6C, Ly6G, F480, PD-L1 and MHC II were evaluated. Then, the same FlowSOM map was applied to WT and SCID groups. In this analysis, visualized cell clusters were distributed in a minimal spanning tree. Each cluster was colored and contained nodes, where the size of each node corresponded with the number of cells represented by the node. These nodes connected to others with the greatest similarities, but the analysis also considered the multidimensional topology of the data. Inside each node, a pie chart indicated the mean fluorescence intensities of the selected markers (49). T-distributed stochastic neighbor embedding (t-SNE) clustering was done based on CD45, NK1.1, CD3, Gr-1, CD11b, CD11c, MHC II, Ly6C, Ly6G, B220, CD4, CD8, PD1, PD-L1, CD25 and F4/80. To determine the enrichment of generated clusters, CD45^+^ cells from WT tumors and SCID tumors were overlaid on the generated t-SNE map. These clusters were further analyzed based on their phenotypic similarities, using PhenoGraph. ClusterExplorer analysis was done based on the median fluorescent intensity of selected surface markers expressed by each of the studied clusters. Here, the expression of CD45, CD3, CD4, CD8, CD25, PD1, Ly6C in T cells was evaluated.

### Flow Cytometry Analysis and FACS Sorting

For myeloid cell analysis, cells were labeled with either a Zombie NIR™ Fixable Viability Kit (BioLegend, #423105) before Fc blocking and cocktail antibody staining, or with Helix NP Green (Biolegend, #425303) after cocktail antibody staining. Cells were washed and incubated for 10 minutes with CD16/CD32 (clone 2.4G2; 553141) to block Fc receptors. Then, cells were washed and a cocktail of antibodies was added: CD11b (Biolegend, clone M1/70 #101207), Ly-6C (Biolegend, clone HK1.4, #128015), Ly-6G (Biolegend, clone 1A8, #127627) with/out CCR1 (Biolegend, clone S15040E, #152505). After staining, cells were acquired with a BD LSRFortessa Cell Analyzer (BD Biosciences) and analyzed with FlowJo, or cells were isolated using a BD FACSAria IIu cytometer.

For CD45.2 staining, cells were stained with Helix NP Blue (Biolegend, #425305) and CD45.2 (Militenyi Biotec, clone 104-2, #130-102-471). After staining, cells were acquired with a BD FACSAria IIu cytometer and analyzed with FlowJo.

## Notes

### Competing Interest Statement

The authors have declared no competing interest.

