## Supplementary Tables and Figures for "Immune Selection Pressure Contributes to Pancreatic Cancer Immune Evasion"

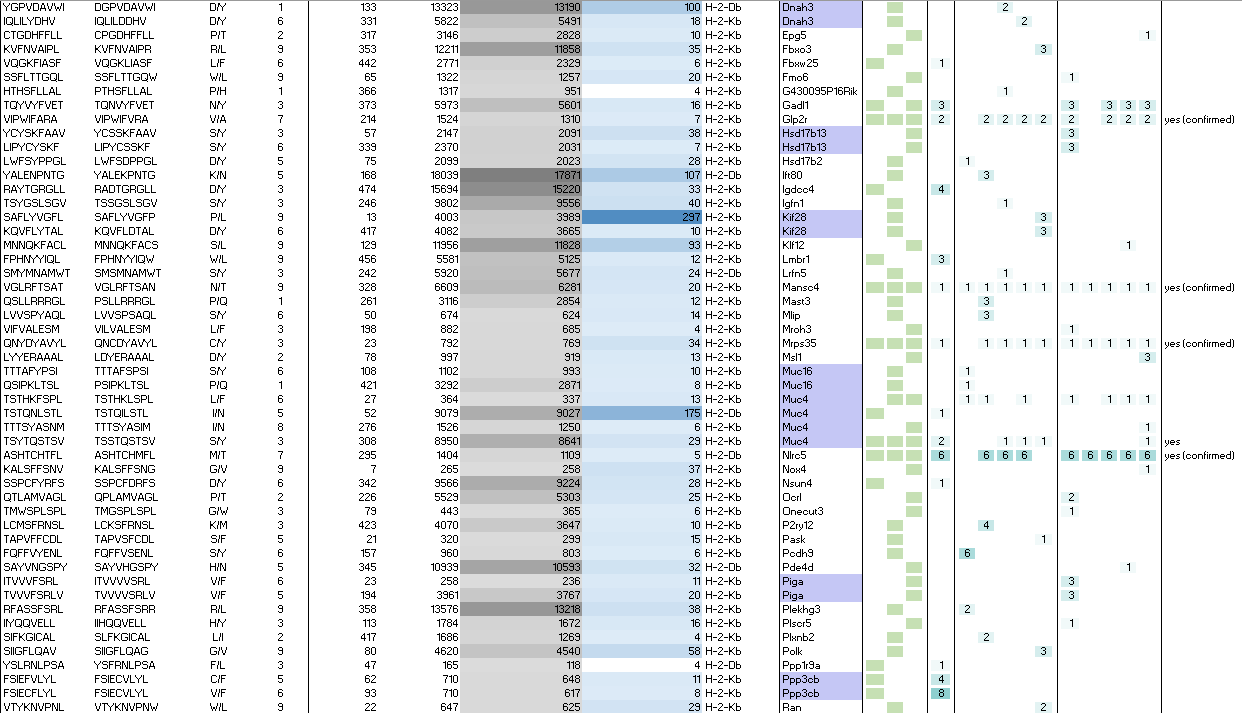

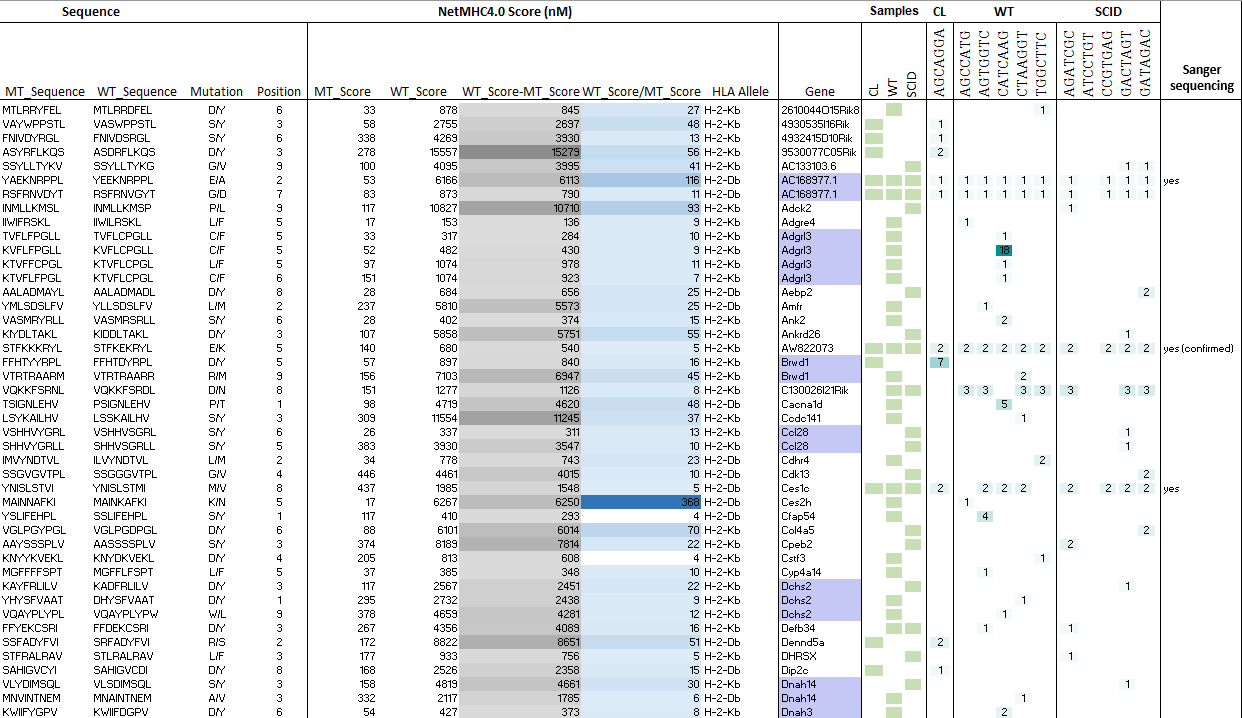
**Supplementary Table 1. Neoepitopes predicted by pVACtools.** The table illustrates the number of putative neoepitopes predicted by pVACtools in the mT3-2D cell lines, 5 WT tumors and 5 SCID tumors.

**Supplementary Table 1 (Cont.). Neoepitopes predicted by pVACtools.**

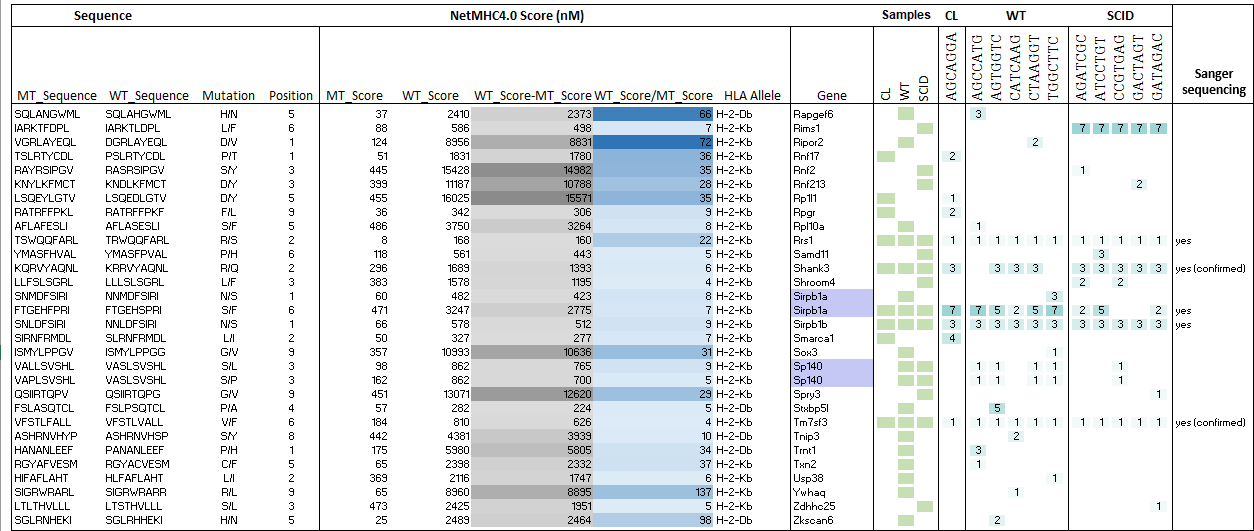

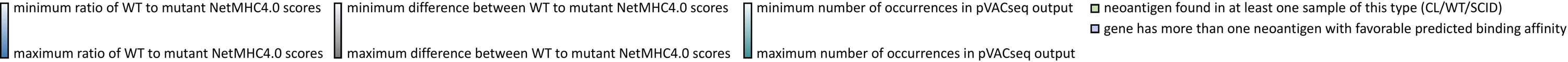

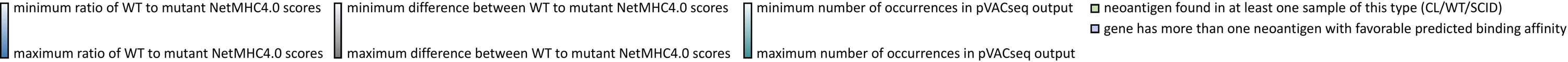

**Supplementary Table 2. Neoepitopes predicted by pVACtools and validated by Sanger Sequencing.** The table illustrates the number of putative neoepitopes shared between the mT3-2D cell line, WT tumors and SCID tumors predicted by pVACtools, and validated by Sanger sequencing in pancreatic tissue from a naïve mouse, mT3-2D cell lines, 3 WT tumors and 3 SCID tumors.

| **Sample ID** | **Gene** | **rsID** | **Alleles** | **WT Flanking Sequence** |
| --- | --- | --- | --- | --- |
| Pancreatic Tissue | Shank3 | N/A | G/G | CGATACAAGCAGAGAGTTTAT |
| mT3-2D cell line | Shank3 | N/A | A/G | CGATACAAGCAGAGAGTTTAT |
| WT tumor | Shank3 | N/A | G/G | CGATACAAGCAGAGAGTTTAT |
| WT tumor | Shank3 | N/A | G/G | CGATACAAGCAGAGAGTTTAT |
| WT tumor | Shank3 | N/A | A/G | CGATACAAGCAGAGAGTTTAT |
| SCID tumor | Shank3 | N/A | A/G | CGATACAAGCAGAGAGTTTAT |
| SCID tumor | Shank3 | N/A | A/G | CGATACAAGCAGAGAGTTTAT |
| SCID tumor | Shank3 | N/A | A/G | CGATACAAGCAGAGAGTTTAT |
| Pancreatic Tissue | Tm7sf3 | rs50547905 | G/G | CTCCACTTTGGTTGCCCTGCT |
| mT3-2D cell line | Tm7sf3 | rs50547905 | G/T | CTCCACTTTGGTTGCCCTGCT |
| WT tumor | Tm7sf3 | rs50547905 | G/T | CTCCACTTTGGTTGCCCTGCT |
| WT tumor | Tm7sf3 | rs50547905 | G/T | CTCCACTTTGGTTGCCCTGCT |
| WT tumor | Tm7sf3 | rs50547905 | G/T | CTCCACTTTGGTTGCCCTGCT |
| SCID tumor | Tm7sf3 | rs50547905 | G/T | CTCCACTTTGGTTGCCCTGCT |
| SCID tumor | Tm7sf3 | rs50547905 | G/T | CTCCACTTTGGTTGCCCTGCT |
| SCID tumor | Tm7sf3 | rs50547905 | G/T | CTCCACTTTGGTTGCCCTGCT |
| Pancreatic Tissue | AW822073 | rs1135301148 | T/T | TATCTCTTCTCCTTAAAAGTT |
| mT3-2D cell line | AW822073 | rs1135301148 | C/C | TATCTCTTCTCCTTAAAAGTT |
| WT tumor | AW822073 | rs1135301148 | C/T | TATCTCTTCTCCTTAAAAGTT |
| WT tumor | AW822073 | rs1135301148 | C/T | TATCTCTTCTCCTTAAAAGTT |
| WT tumor | AW822073 | rs1135301148 | C/T | TATCTCTTCTCCTTAAAAGTT |
| SCID tumor | AW822073 | rs1135301148 | C/T | TATCTCTTCTCCTTAAAAGTT |
| SCID tumor | AW822073 | rs1135301148 | C/T | TATCTCTTCTCCTTAAAAGTT |
| SCID tumor | AW822073 | rs1135301148 | C/T | TATCTCTTCTCCTTAAAAGTT |

**Supplementary Table 2 (Cont.). Neoepitopes predicted by pVACtools and validated by Sanger sequencing.**

| **Sample ID** | **Gene** | **rsID** | **Alleles** | **WT Flanking Sequence** |
| --- | --- | --- | --- | --- |
| Pancreatic Tissue | Glp2r | rs28217197 | T/T | TGGATTTTTGTCCGAGCAAGC |
| mT3-2D cell line | Glp2r | rs28217197 | C/C | TGGATTTTTGTCCGAGCAAGC |
| WT tumor | Glp2r | rs28217197 | T/T | TGGATTTTTGTCCGAGCAAGC |
| WT tumor | Glp2r | rs28217197 | T/T | TGGATTTTTGTCCGAGCAAGC |
| WT tumor | Glp2r | rs28217197 | T/C | TGGATTTTTGTCCGAGCAAGC |
| SCID tumor | Glp2r | rs28217197 | T/C | TGGATTTTTGTCCGAGCAAGC |
| SCID tumor | Glp2r | rs28217197 | T/T | TGGATTTTTGTCCGAGCAAGC |
| SCID tumor | Glp2r | rs28217197 | T/T | TGGATTTTTGTCCGAGCAAGC |
| Pancreatic Tissue | Mansc4 | rs51447380 | A/A | ACATCTGCAAACGTGTCCACT |
| mT3-2D cell line | Mansc4 | rs51447380 | A/C | ACATCTGCAAACGTGTCCACT |
| WT tumor | Mansc4 | rs51447380 | A/C | ACATCTGCAAACGTGTCCACT |
| WT tumor | Mansc4 | rs51447380 | A/C | ACATCTGCAAACGTGTCCACT |
| WT tumor | Mansc4 | rs51447380 | A/C | ACATCTGCAAACGTGTCCACT |
| SCID tumor | Mansc4 | rs51447380 | A/C | ACATCTGCAAACGTGTCCACT |
| SCID tumor | Mansc4 | rs51447380 | A/C | ACATCTGCAAACGTGTCCACT |
| SCID tumor | Mansc4 | rs51447380 | A/C | ACATCTGCAAACGTGTCCACT |
| Pancreatic Tissue | Mrps35 | rs36949762 | G/G | CGGCAGAACTGTGACTATGCA |
| mT3-2D cell line | Mrps35 | rs36949762 | G/A | CGGCAGAACTGTGACTATGCA |
| WT tumor | Mrps35 | rs36949762 | G/G | CGGCAGAACTGTGACTATGCA |
| WT tumor | Mrps35 | rs36949762 | G/A | CGGCAGAACTGTGACTATGCA |
| WT tumor | Mrps35 | rs36949762 | G/A | CGGCAGAACTGTGACTATGCA |
| SCID tumor | Mrps35 | rs36949762 | G/A | CGGCAGAACTGTGACTATGCA |
| SCID tumor | Mrps35 | rs36949762 | G/A | CGGCAGAACTGTGACTATGCA |
| SCID tumor | Mrps35 | rs36949762 | G/A | CGGCAGAACTGTGACTATGCA |
| Pancreatic Tissue | Nlrc5 | rs32684635 | T/T | ACCTGCCACATGTTCCTCTGC |
| mT3-2D cell line | Nlrc5 | rs32684635 | C/T | ACCTGCCACATGTTCCTCTGC |
| WT tumor | Nlrc5 | rs32684635 | T/T | ACCTGCCACATGTTCCTCTGC |
| WT tumor | Nlrc5 | rs32684635 | T/T | ACCTGCCACATGTTCCTCTGC |
| WT tumor | Nlrc5 | rs32684635 | C/T | ACCTGCCACATGTTCCTCTGC |
| SCID tumor | Nlrc5 | rs32684635 | C/T | ACCTGCCACATGTTCCTCTGC |
| SCID tumor | Nlrc5 | rs32684635 | C/T | ACCTGCCACATGTTCCTCTGC |
| SCID tumor | Nlrc5 | rs32684635 | C/T | ACCTGCCACATGTTCCTCTGC |

**Supplementary Table 3. Differentially expressed genes between the WT and SCID malignant epithelial pancreatic cells.** The table illustrates the 52 genes that were differentially expressed between WT and SCID malignant cells using RNA-seq. (Ch.=chromosome).

| Ch. | Start | Stop | Strand | Transcript | Gene.Symbol | SCID vs. Cell (Log2 FC) | WT vs. Cell (Log2 FC) | SCID vs. WT  (Log2 FC) |
| --- | --- | --- | --- | --- | --- | --- | --- | --- |
| 11 | 83572917 | 83578637 | - | NM_011338 | **Ccl9** | 0.72 | 2.60 | -1.88 |
| 17 | 34153191 | 34160230 | + | NM_010387 | H2-DMb1 | 0.40 | 2.20 | -1.80 |
| X | 20925454 | 20931556 | - | NM_008823 | Cfp | 0.93 | 2.71 | -1.78 |
| 3 | 87250965 | 87263525 | - | NM_030707 | Fcrls | 0.65 | 2.42 | -1.77 |
| 7 | 19696244 | 19699189 | - | NM_009696 | Apoe | 1.02 | 2.77 | -1.75 |
| 7 | 100937634 | 100964367 | - | NM_183168 | P2ry6 | 0.18 | 1.74 | -1.56 |
| 14 | 70774381 | 70778495 | + | NM_010071 | Dok2 | 0.37 | 1.92 | -1.54 |
| 10 | 24915207 | 24927471 | - | NM_007482 | **Arg1** | 0.72 | 2.22 | -1.50 |
| 19 | 6998958 | 7019470 | - | NM_153795 | Fermt3 | 0.72 | 2.20 | -1.49 |
| 9 | 120048683 | 120068297 | - | NM_009987 | Cx3cr1 | 0.59 | 2.06 | -1.47 |
| 2 | 131992850 | 132029989 | - | NM_175445 | Rassf2 | 0.82 | 2.22 | -1.41 |
| 19 | 6844623 | 6858212 | - | NM_001081291 | Ccdc88b | 0.61 | 2.01 | -1.40 |
| 3 | 83836272 | 83841609 | - | NM_011905 | Tlr2 | 0.77 | 2.17 | -1.40 |
| 17 | 71252176 | 71310966 | - | NM_145158 | Emilin2 | 1.30 | 2.64 | -1.34 |
| 2 | 44983512 | 45113280 | - | NM_015753 | Zeb2 | 0.87 | 2.19 | -1.32 |
| 19 | 61224402 | 61228419 | - | NM_009970 | Csf2ra | 0.75 | 2.06 | -1.32 |
| 6 | 97286867 | 97487821 | - | NM_001346637 | Frmd4b | -0.48 | 0.83 | -1.31 |
| 6 | 122847140 | 122856158 | - | NM_009779 | C3ar1 | -0.10 | 1.20 | -1.30 |
| 12 | 103442166 | 103443681 | - | NM_029803 | Ifi27l2a | 0.20 | 1.48 | -1.27 |
| 11 | 69664213 | 69666171 | - | NM_001291058 | **Cd68** | 0.99 | 2.25 | -1.26 |
| 8 | 68880555 | 68906933 | + | NM_008509 | Lpl | -0.78 | 0.48 | -1.26 |
| 19 | 3935186 | 3949341 | + | NM_019449 | Unc93b1 | 0.62 | 1.87 | -1.26 |
| 1 | 52119622 | 52161866 | + | NM_001205313 | **Stat1** | -0.06 | 1.19 | -1.25 |
| 15 | 6386748 | 6440710 | + | NM_023118 | Dab2 | 0.22 | 1.47 | -1.25 |
| 11 | 58199556 | 58207593 | + | NM_018738 | Igtp | 0.89 | 2.13 | -1.24 |
| 17 | 33996012 | 34000334 | - | NM_001001892 | H2-K1 | -0.24 | 0.99 | -1.24 |

**Supplementary Table 3 (Cont.). Differentially expressed genes between the WT and SCID malignant epithelial pancreatic cells**

| Ch. | Start | Stop | Strand | Transcript | Gene.Symbol | SCID vs. Cell (Log2 FC) | WT vs. Cell (Log2 FC) | SCID vs. WT  (Log2 FC) |
| --- | --- | --- | --- | --- | --- | --- | --- | --- |
| 7 | 29221928 | 29232523 | - | NM_001033525 | Kcnk6 | 0.42 | 1.65 | -1.23 |
| 15 | 6386748 | 6440710 | + | NM_001310446 | Dab2 | 0.24 | 1.47 | -1.23 |
| 6 | 124720707 | 124733155 | - | NM_013545 | Ptpn6 | 0.28 | 1.49 | -1.20 |
| 17 | 33996012 | 34000348 | - | NM_001347346 | H2-K1 | -0.67 | 0.51 | -1.18 |
| 13 | 60602211 | 60763192 | + | NM_134062 | Dapk1 | -0.74 | 0.42 | -1.15 |
| 5 | 105078394 | 105110293 | - | NM_172777 | Gbp9 | 0.65 | 1.80 | -1.15 |
| 13 | 23702034 | 23710855 | - | NM_010424 | Hfe | 0.34 | 1.49 | -1.15 |
| 8 | 117498275 | 117635143 | + | NM_172285 | Plcg2 | 0.38 | 1.49 | -1.11 |
| 7 | 68736994 | 68749239 | - | NM_001042592 | Arrdc4 | 0.84 | 1.95 | -1.11 |
| 17 | 35263094 | 35267498 | + | NM_010380 | H2-D1 | -0.29 | 0.81 | -1.10 |
| 1 | 105780723 | 105847982 | + | NM_009399 | Tnfrsf11a | -0.14 | 0.95 | -1.10 |
| 9 | 88548020 | 88571087 | + | NM_001142943 | Zfp949 | -0.57 | 0.52 | -1.09 |
| 5 | 100679484 | 100719717 | - | NM_152803 | Hpse | -0.41 | 0.68 | -1.09 |
| 7 | 30423304 | 30428747 | + | NM_172142 | Nfkbid | 0.23 | 1.30 | -1.07 |
| 17 | 8128591 | 8147833 | - | NM_026611.1 | Rnaset2b | -0.69 | 0.36 | -1.05 |
| 11 | 53770473 | 53777325 | + | NM_001159396 | Irf1 | -0.06 | 0.99 | -1.04 |
| 17 | 46630629 | 46646081 | - | NM_001357130 | Klc4 | -0.43 | 0.60 | -1.04 |
| 16 | 35832878 | 35871383 | - | NM_001039530 | Parp14 | 1.33 | 2.36 | -1.03 |
| 8 | 70762773 | 70766664 | - | NM_023065 | Ifi30 | -0.23 | 0.80 | -1.03 |
| 2 | 125830302 | 125859083 | - | NM_001285513 | Cops2 | -0.79 | 0.23 | -1.02 |
| 5 | 43818893 | 43843469 | + | NM_009763 | Bst1 | 0.13 | 1.14 | -1.01 |
| X | 107397195 | 107403361 | - | NM_008409 | Itm2a | 1.38 | 0.36 | 1.02 |
| Y | 1260715 | 1286614 | - | NM_012008 | Ddx3y | 0.31 | -0.71 | 1.02 |
| 16 | 13833573 | 13903146 | - | NM_001039533 | Pdxdc1 | 0.60 | -0.46 | 1.06 |
| 17 | 66078795 | 66101560 | - | NM_028388 | Ndufv2 | -0.74 | -1.85 | 1.11 |
| 6 | 145211147 | 145216543 | + | NM_133688 | Etfrf1 | 0.42 | -0.72 | 1.14 |

**Supplementary Table 4. Hallmark pathway analysis of differentially expressed genes between the WT and SCID malignant epithelial pancreatic cells.** The table illustrates the Hallmark pathways that were significantly different between WT and SCID malignant cells using RNA-seq. Green arrows indicate pathways that were upregulated in WT malignant cells compared to SCID malignant cells. Red arrows indicate pathways that were downregulated in WT malignant cells compared to SCID malignant cells.

| Hallmark pathways | WT vs.  SCID | Adjusted  p-values |
| --- | --- | --- |
| HALLMARK_INTERFERON_GAMMA_RESPONSE |  | 1.72E-11 |
| HALLMARK_INTERFERON_ALPHA_RESPONSE |  | 1.29E-06 |
| HALLMARK_IL6_JAK_STAT3_SIGNALING |  | 0.000288205 |
| HALLMARK_XENOBIOTIC_METABOLISM |  | 0.003418453 |
| HALLMARK_COMPLEMENT |  | 0.023539937 |
| HALLMARK_ALLOGRAFT_REJECTION |  | 0.023539937 |
| HALLMARK_EPITHELIAL_MESENCHYMAL_TRANSITION |  | 0.000407251 |
| HALLMARK_HYPOXIA |  | 0.007250325 |

**Supplementary Table 5. Western blot antibodies.** The table describes the antibodies used for Western blot studies.

| Proteins | Primary Antibodies | Secondary Antibodies |
| --- | --- | --- |
| Tubulin | Abcam, #ab6160  diluted 1:1000 | Anti-rat IgG: Cell Signaling, #7077  diluted 1:1000 |
| GAPDH | Cell Signaling, #5174  diluted 1:5000 | Anti-rabbit IgG: Cell Signaling, #7074  diluted 1:2000 |
| S100a9 | Abcam, #ab105472  diluted 1:1000 | Anti-rabbit IgG: Cell Signaling, #7074  diluted 1:1000 |
| Arginase1 | Abcam, #ab60176  diluted 1:1000 | Anti-goat IgG: Jackson ImmunoResearch, #705-035-147  diluted 1:00 |
| STAT1 | Cell Signaling, #14994S  diluted 1:1000 | Anti-rabbit IgG: Cell Signaling, #7074  diluted 1:1000 |
| CCL9 | For **Figure 5b**:  R&D Systems, # MAB463  diluted 1:1000 | Anti-rat IgG: Cell Signaling, #7077  diluted 1:1000 |
|  | For **Supplementary Figure 5b**:  Abcam, #ab9913  diluted 1:500 | Anti-rabbit IgG: Cell Signaling, #7074  diluted 1:1000 |

**Supplementary Table 6. RT-qPCR primer sequences.** The table describes the primer sequences used for RT-qPCR studies.

| Genes | Forward Primers | Reverse Primers |
| --- | --- | --- |
| *Hprt* | GAT TAG CGA TGA TGA ACC AGG TT | CCT CCC ATC TCC TTC ATG ACA |
| *Ifng* | GGA TGC ATT CAT GAG TAT TGC | GTG GAC CAC TCG GAT GAG |
| *Ptprc* | CCA CCA GGG ACT GAC AAG TT | TGT AAT TTG TTT GGG CAC GA |
| *S100a8* | CCG TCT TCA AGA CAT CGT TTG A | GTA GAG GGC ATG GTG ATT TCC T |
| *S100a9* | GTT GAT CTT TGC CTG TCA TGA G | AGC CAT TCC CTT TAG ACT TGG |
| *Stat1* | CTG AAT ATT TCC CTC CTG GG | TCC CGT ACA GAT GTC CAT GAT |
| *Ccl9* | TGG GCC CAG ATC ACA CAT GCA AC | CGG CCT GGT ACA CCC ACC AC |
| *Arg1* | GGA ATC TGC ATG GGC AAC CTG TGT | AGG GTC TAC GTC TCG CAA GCCA |
| *Cd68* | CTG TTT GGC TCT GCC ATA GGA G | CCT AGG ATG GCA TTT GTT GAT GTG G |
| *Gapdh* | TCA CCA CCA TGG AGA AGG C | GCT AAG CAG TTG GTG GTG CA |

**Supplementary Table 7. Immunohistochemistry antibodies.** The table describes the antibodies used for immunohistochemistry studies.

| Proteins | Primary Antibodies |
| --- | --- |
| Ki67 | BIOCARE, #CRM325, diluted 1:80 |
| CD4 | Cell Signaling, #25229, diluted 1:25 |
| CD8 | Cell Signaling, #98941, diluted 1:25 |
| CD68 | Thermo Scientific, #PA5-78996 |
| S100a8 | Abcam, #ab92331, diluted 1:1000 |
| S100a9 | Cell Signaling, #73425, diluted 1:1000 |

**Supplementary Figures**

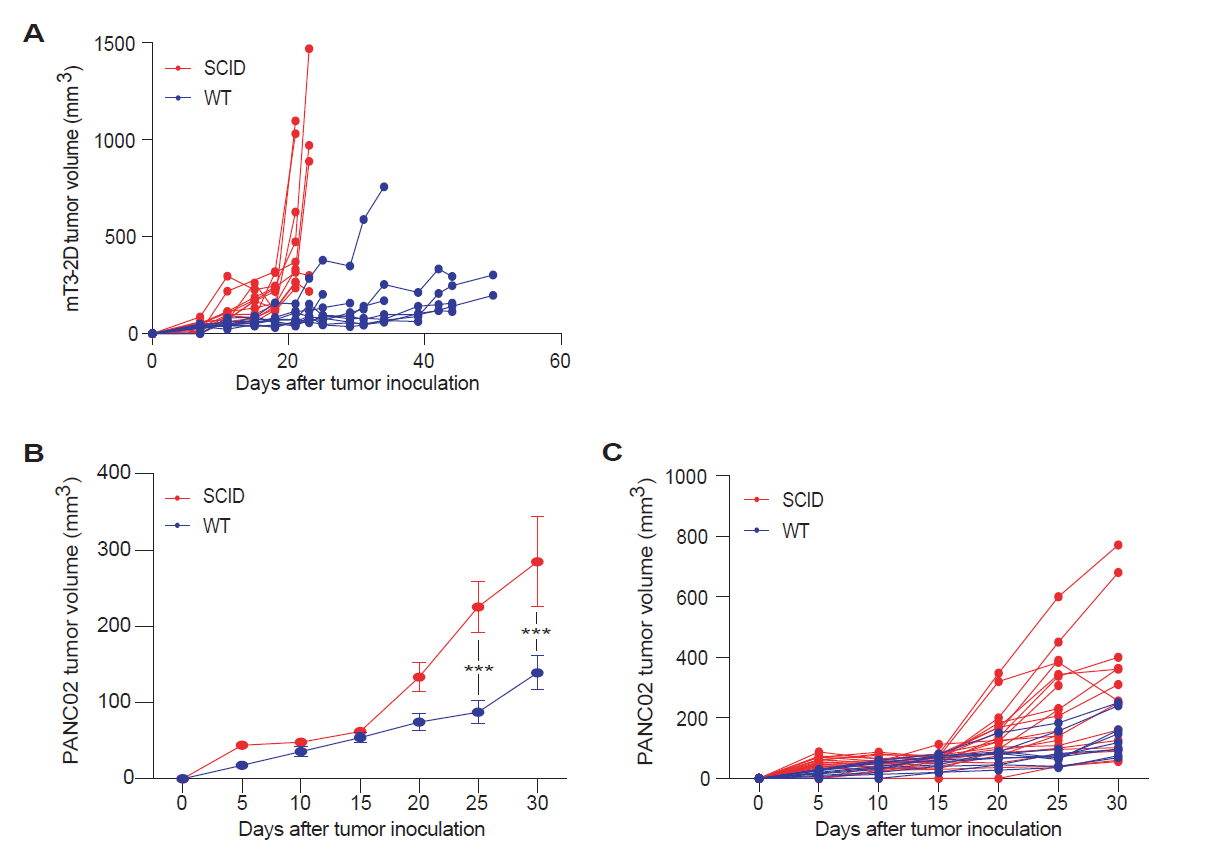

**Supplementary Figure 1. T cell immunity incompletely regulates pancreatic cancer tumor growth. (A)** Growth curves depicting individual mT3-2D tumors grown in syngeneic WT (n=9) and SCID (n=13) C57BL/6J mice. **(B)** Growth curves depicting average PANC02 tumor volumes (±SEM) grown in syngeneic WT (n=10) and SCID (n=20) C57BL/6J mice. **(C)** Growth curves depicting individual PANC02 tumors grown in syngeneic WT (n=10) and SCID (n=20) C57BL/6J mice.

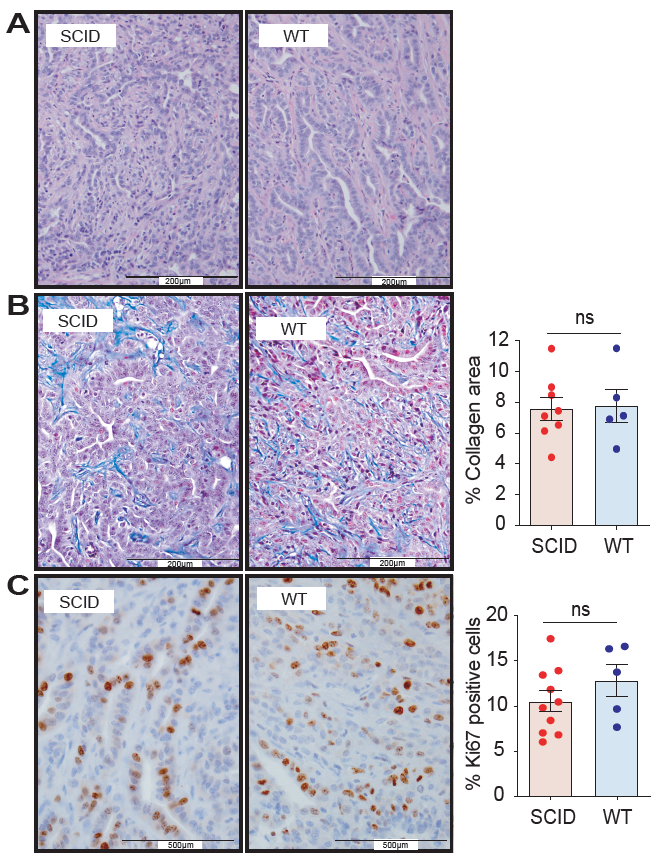

**Supplementary Figure 2. Histology of mT3-2D subcutaneous tumors grown in immunocompetent and immunodeficient mice. (A)** Representative images of mT3-2D tumor tissues from WT (n=5) and SCID (n=8) mice subjected to hematoxylin and eosin (H&E) staining. **(B)** Representative images of mT3-2D tumor tissues from WT (n=5) and SCID (n=8) mice subjected to Masson’s trichrome staining, which stains collagen blue. Quantification of Masson’s trichrome-stained collagen expression using imageJ is shown in the bar plot to the right. **(C)** Representative images of mT3-2D tumor tissues from WT (n=5) and SCID (n=10) mice subjected to Ki67 immunohistochemistry (IHC) staining. Quantification of Ki67 positive cells using FIJI imageJ is shown in the bar plot to the right. Data represented as mean (±SEM) and a Mann-Whitney test was used, (ns=not significant).

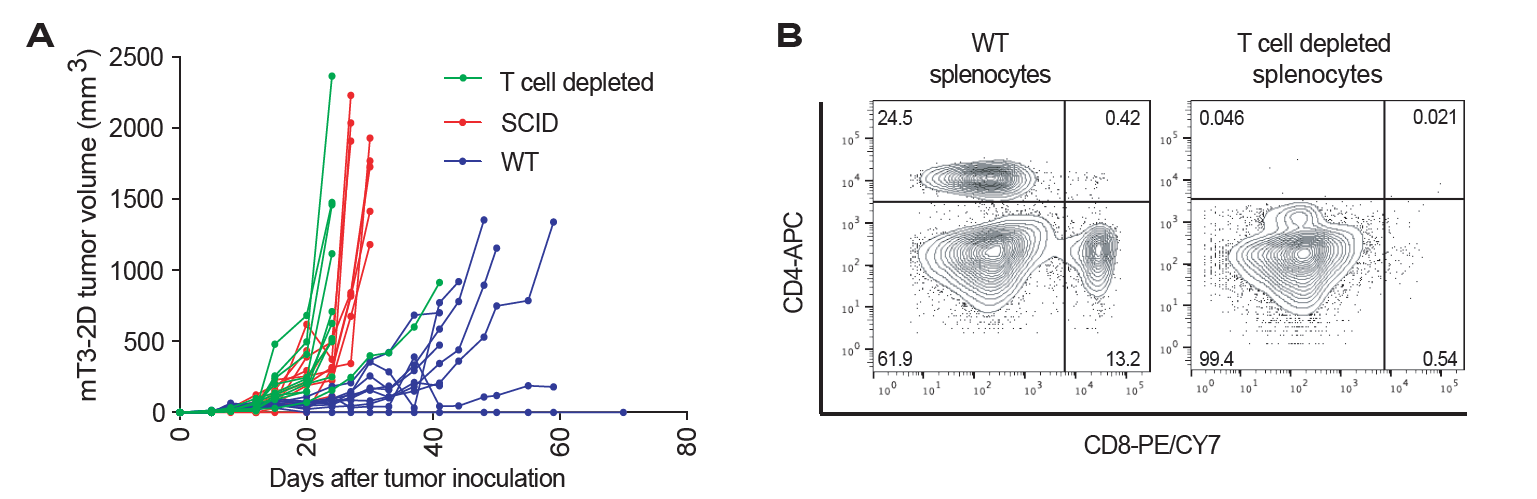

**Supplementary Figure 3. Delayed mT3-2D tumor growth in immunocompetent mice is T cell dependent. (A)** Growth curves depicting individual mT3-2D tumors grown in syngeneic WT, SCID, and T cell depleted WT C57BL/6J mice (n=10 per group). **(B)** The efficiency of *in vivo* T cell depletion was determined using flow cytometry analysis of splenocytes from T cell depleted mice compared to control mice after 1 week of treatment (2 doses).

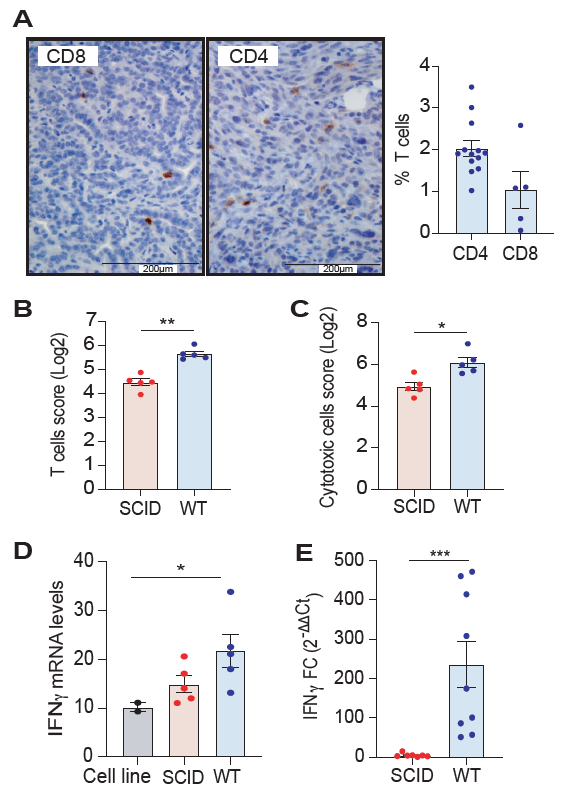

**Supplementary Figure 4. T cell immunity is induced in mT3-2D WT tumors. (A)** Quantification of CD4 and CD8 positive cells using FIJI imageJ is shown in the bar plot to the right. Data represented as mean (±SEM) and Mann-Whitney test was used. **(B)** “T cell” score and **(C)** “cytotoxic cell” score analyzed using the NanoString nSolver tool (n=5 per group). Data represented as mean (±SEM) and a Mann-Whitney test was used. **(D)** *Ifng* RNA counts as determined using the NanoString nSolver tool (n=2 technical replicates for the cell line, n=5 biological replicates for the tumor samples per group). Data represented as mean (±SEM) and a Kruskal-Wallis test was used. **(E)** Quantitative real-time PCR analysis of *Ifng* gene expression in WT (n=9) and SCID tumors (n=7). Data represented as mean FC (fold change) (±SEM) and a Mann-Whitney test was used. One asterisk (*) indicates p value < 0.05, 2 asterisks (**) indicates p value < 0.01 and 3 asterisks (***) indicates p value < 0.005.

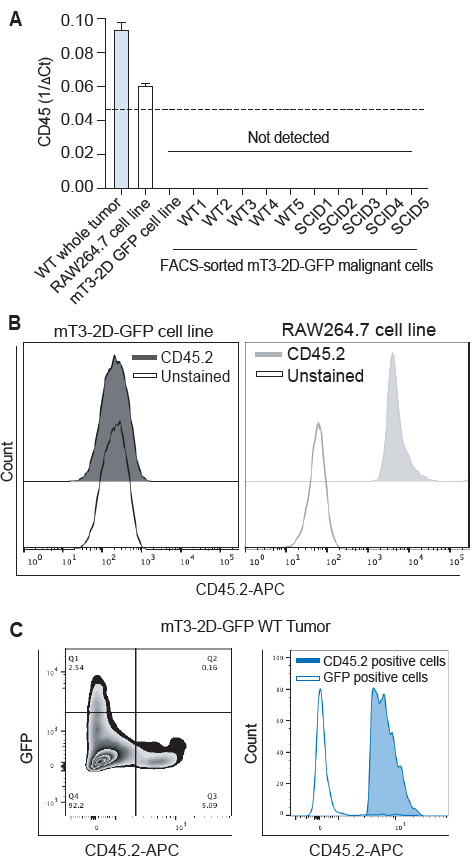

**Supplementary Figure 5. FACS-sorted mT3-2D GFP cells and the mT3-2D cell line do not express CD45. (A)** Quantitative real-time PCR analysis of *Ptprc* expression in a whole WT tumor (including the stroma as a positive control), RAW264.7 cell line (positive control), mT3-2D cell line, and FACS-sorted mT3-2D GFP cells from WT and SCID tumors. Data represented as mean (±SEM), and the dashed line indicates the detection threshold. **(B)** Flow cytometry analysis of CD45.2 in mT3-2D GFP cell line and RAW264.7 cell line (positive control) analyzed using FlowJo**. (C)** Flow cytometry analysis of CD45.2 and GFP in a WT mT3-2D-GFP tumor, analyzed using FlowJo. CD45.2 fluorescence intensity in GFP positive cells (Q1) and CD45.2 positive cells (Q3) is shown in the histogram to the right.

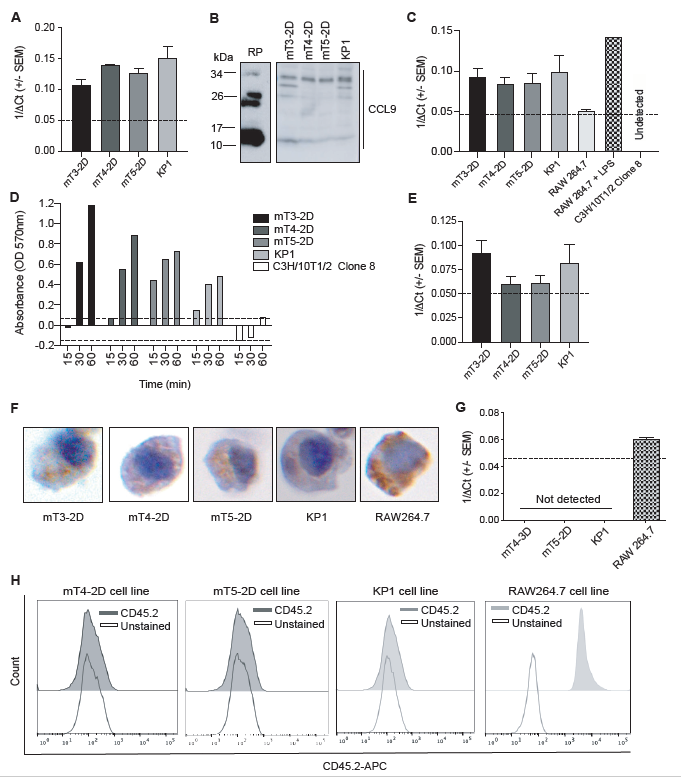

**Supplementary Figure 6. KPC cell lines express basal level of myeloid markers but do not express CD45. (A)** Quantitative real-time PCR analysis of *Ccl9* expression in KPC cell lines. Data represented as mean (±SEM), and the dashed line indicates the detection threshold. **(B)** Western blot of CCL9 in recombinant protein (RP) (Novus Biologicals, NBP2-35086) and KPC cell lines. **(C)** Quantitative real-time PCR analysis of *Arg1* expression in KPC cell lines, RAW264.7 cell line, LPS-treated RAW264.7 cells (positive control) and C3H/10T1/2 Clone 8 cell line (negative control). Data represented as mean (±SEM), and the dashed line indicates the detection threshold. **(D)** Arginase activity in KPC cell lines, and C3H/10T1/2 Clone 8 cell line (negative control), using arginase activity assay and determined by the absorbance of H_2_O_2_ at the indicated time points. Data represented as mean (±SEM), and the dashed line indicates the detection threshold. **(E)** Quantitative real-time PCR analysis of *Cd68* expression in KPC cell lines. Data represented as mean (±SEM), and the dashed line indicates the detection threshold. **(F)** Representative HistoGel images of KPC cells subjected to CD68 IHC staining. **(G)** Quantitative real-time PCR analysis of *Ptprc* expression in KPC cell lines. Data represented as mean (±SEM), and the dashed line indicates the detection threshold. **(H)** Flow cytometry analysis of CD45.2 in KPC cell lines and RAW264.7 cell line (positive control) analyzed using FlowJo.

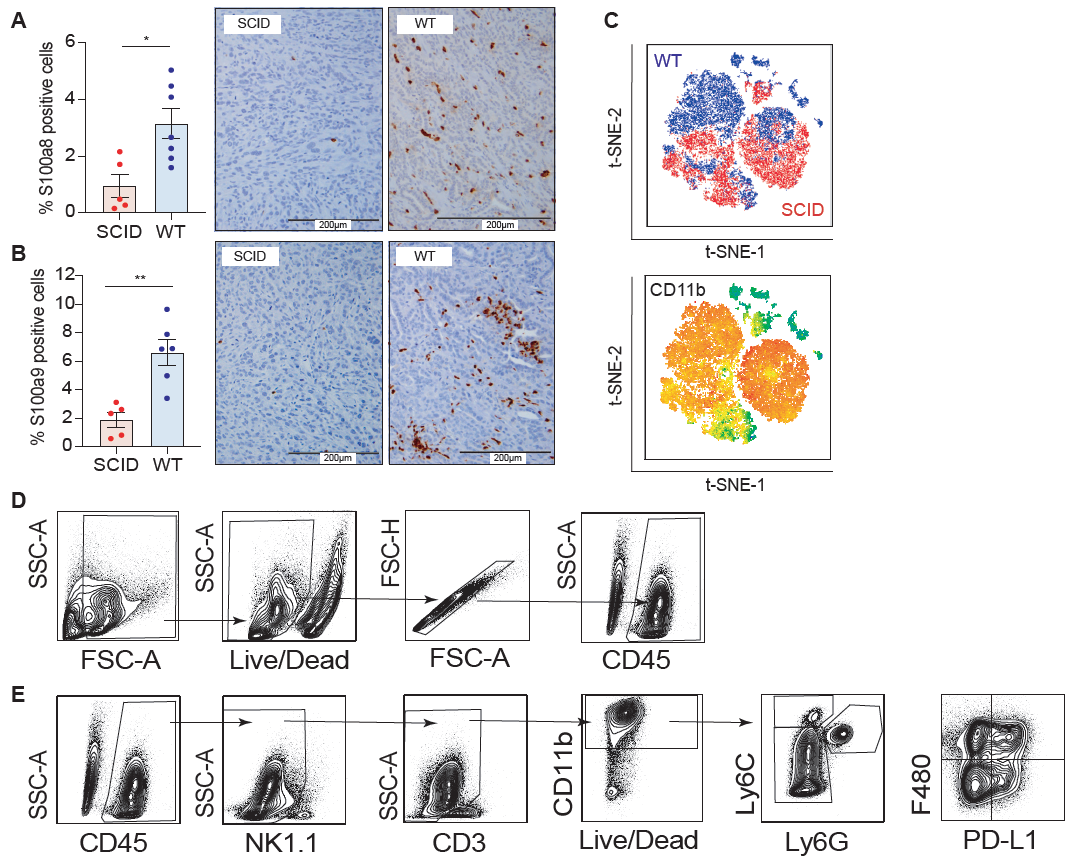

**Supplementary Figure 7. WT and SCID tumors are highly infiltrated by CD11b^+^ myeloid cells. (A-B)** Representative images of mT3-2D tumor tissues from WT (n=7) and SCID (n=5) mice subjected to S100a8 and S100a9 IHC staining, respectively. Quantification of positive cells using FIJI imageJ is shown in the bar plots to the left. Data represented as mean (±SEM) and Mann-Whitney test was used. One asterisk (*) indicates p value < 0.05, and two asterisks (**) indicate p value < 0.01. **(C)** Top: t-SNE plot of immune cells from WT (blue) and SCID (red) tumors. Bottom: the same t-SNE plot visualized based on the expression of CD11b marker, where orange indicates high expression and green indicates low expression. Samples were gated on Live/CD45^+^ cells (n=5 per group). **(D-E)** Flow cytometry gating strategies for myeloid cell analysis. **(D)** Cells were gated based on their size and granularity (SSC-A and FSC-A, respectively). Dead cells and doublets were excluded and CD45^+^ cells were selected. **(E)** CD45^+^ cells were evaluated for the expression of NK1.1 and CD3. Then, C11b^+^ NK1.1^-^/CD3^-^ cells were selected. This population was then divided into three clusters based on their expression of Ly6C and Ly6G. Subsequently, the expression of F480 and PD-L1 in the three clusters was evaluated.

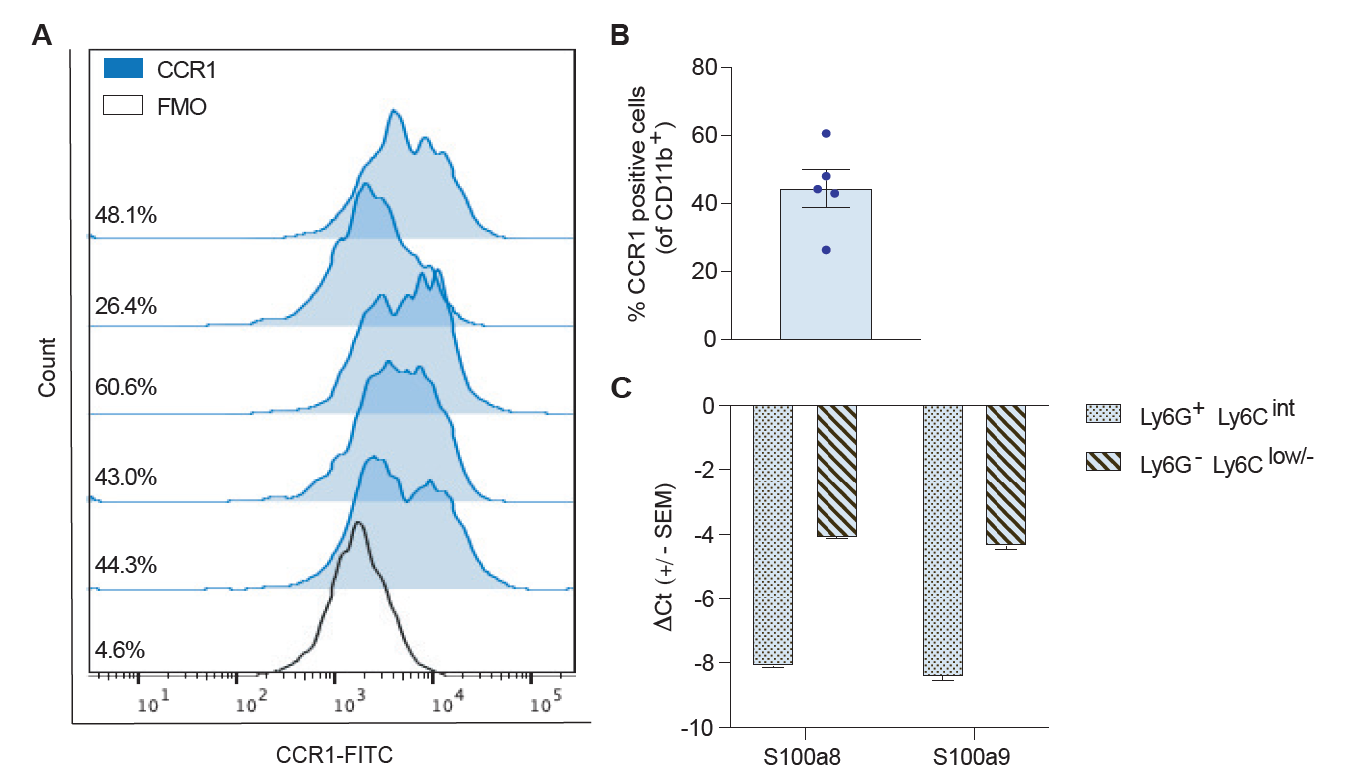

**Supplementary Figure 8. CD11b^+^ myeloid cell populations express CCR1, S100a8 and S100a9. (A)** Flow cytometry analysis of CD11b^+^ cells from five WT tumor samples stained with CCR1, and CCR1 unstained CD11b^+^ cells from combined WT tumor samples (Fluorescence minus one (FMO)). **(B)** Bar chart indicating the percentage of CCR1 positive cells from **(A)**. Data represented as mean (±SEM). **(C)** Quantitative real-time PCR analysis of *S100a8 and S100a9* expression in CD11b^+^ Ly6G^-^ Ly6C^low/-^ and CD11b^+^ Ly6G^+^ splenocytes from naïve mice. Two splenocytes were combined and processed as a single sample. Data represented as mean (±SEM).

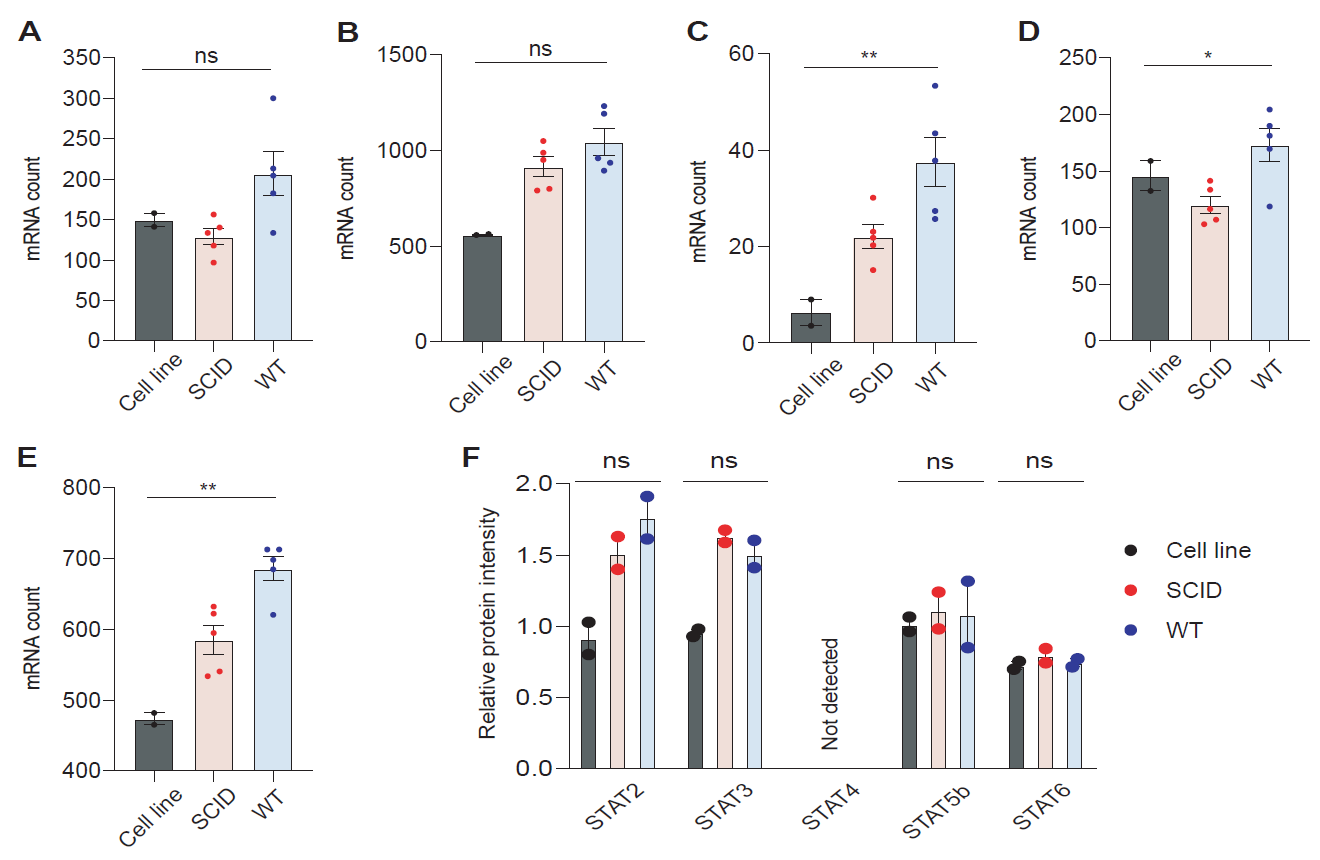

**Supplementary Figure 9. Less notable difference in the expression of STAT genes other than STAT1 between WT and SCID whole tumors.** **(A)** *Stat2* **(B)** *Stat3* **(C)** *Stat4* **(D)** *Stat5a* **(E)** *Stat6* mRNA counts by NanoString nCounter analysis (for the cell line, n=2 technical replicates and for tumor samples, n=5 biological replicates per group). **(F)** Relative protein intensities detected by mass spectrometry (n=2 technical replicates per group). Data represented as mean (±SEM) and a Mann-Whitney test was used. One asterisk (*) indicates p value < 0.05, 2 asterisks (**) indicate p value < 0.01 and ns= not significant.

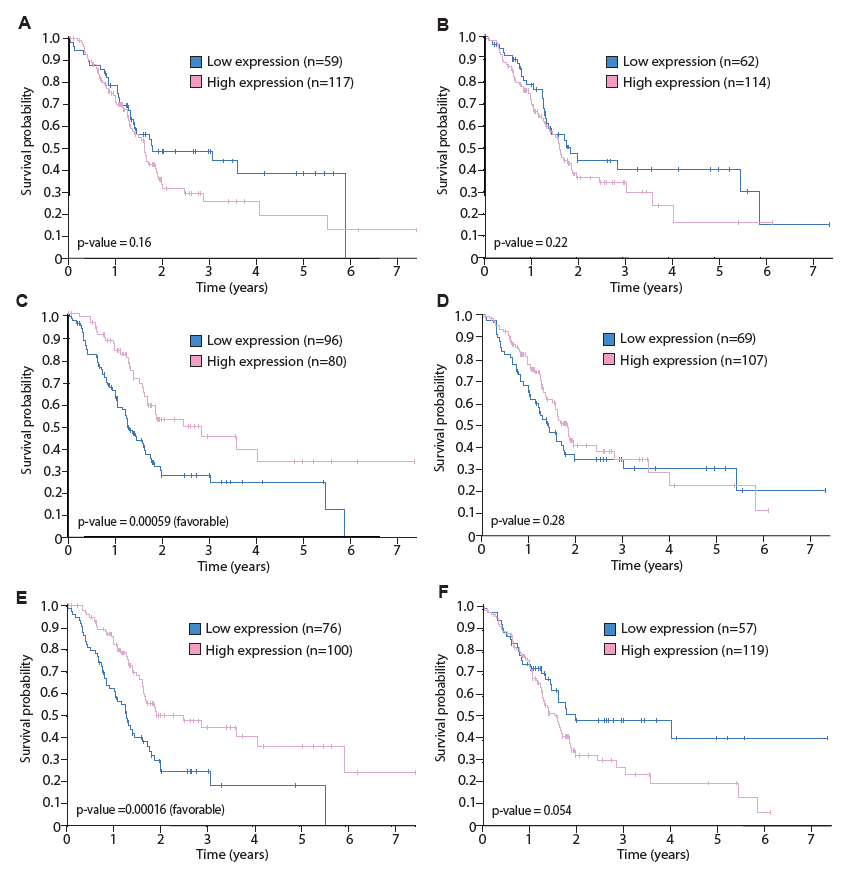

**Supplementary Figure 10. *STAT* genes other than *STAT1* are not significantly associated with shorter overall survival for pancreatic cancer patients.** Kaplan-Meier plots show **(A)** *STAT2* **(B)** *STAT3* **(C)** *STAT4* **(D)** *STAT5A* **(E)** *STAT5B* and **(F)** *STAT6* pancreatic cancer survival analysis using the TCGA data from the Human Protein Atlas. The x-axis shows time for survival (years) and y-axis shows the probability of survival, where 1.0 corresponds to 100 percent.

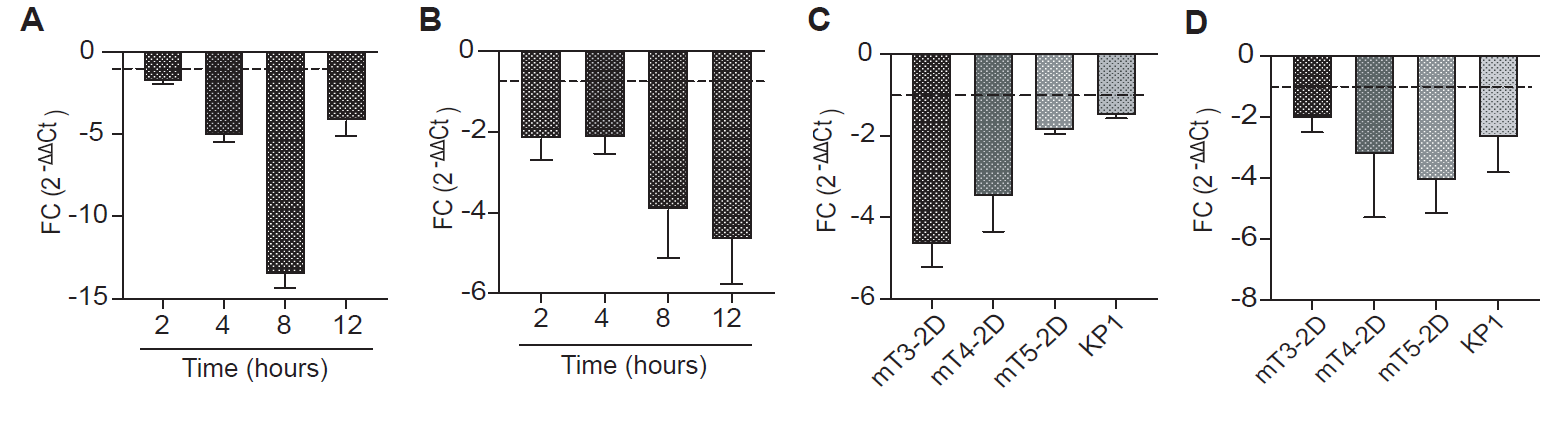

**Supplementary Figure 11. Ruxolitinib downregulates *Ccl9* and *Arg1* expression in murine KPC cell lines *in vitro*. (A)** Quantitative real-time PCR analysis of *Ccl9* expression in mT3-2D cells treated with 1 µM ruxolitinib relative to untreated cells at the indicated time points. **(B)** Quantitative real-time PCR analysis of *Arg1* expression in mT3-2D cells treated with 1 µM ruxolitinib relative to untreated cells at the indicated time points. **(C)** Quantitative real-time PCR analysis of *Ccl9* expression in murine KPC cell lines treated with 1 µM ruxolitinib for 4 hours relative to untreated cells. (D) Quantitative real-time PCR analysis of *Arg1* expression in murine KPC cell lines treated with 1 µM ruxolitinib for 4 hours relative to untreated cells. Data represented as mean FC (fold change) (±SEM).

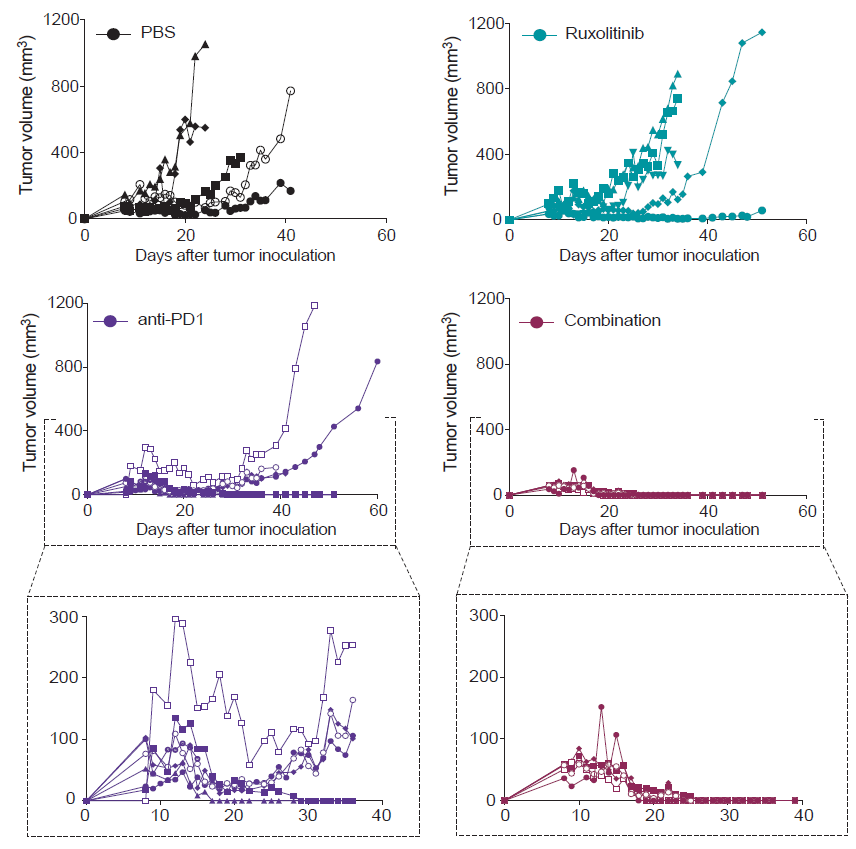

**Supplementary Figure 12. Ruxolitinib improves the anti-tumor efficacy of anti-PD1 treatment.** Individual tumor growth curves of mT3-2D tumors grown in C57BL/6J immunocompetent mice treated with PBS (n=5), ruxolitinib (n=5), anti-PD1 (n=7) or combination of anti-PD1 and ruxolitinib (n=5).

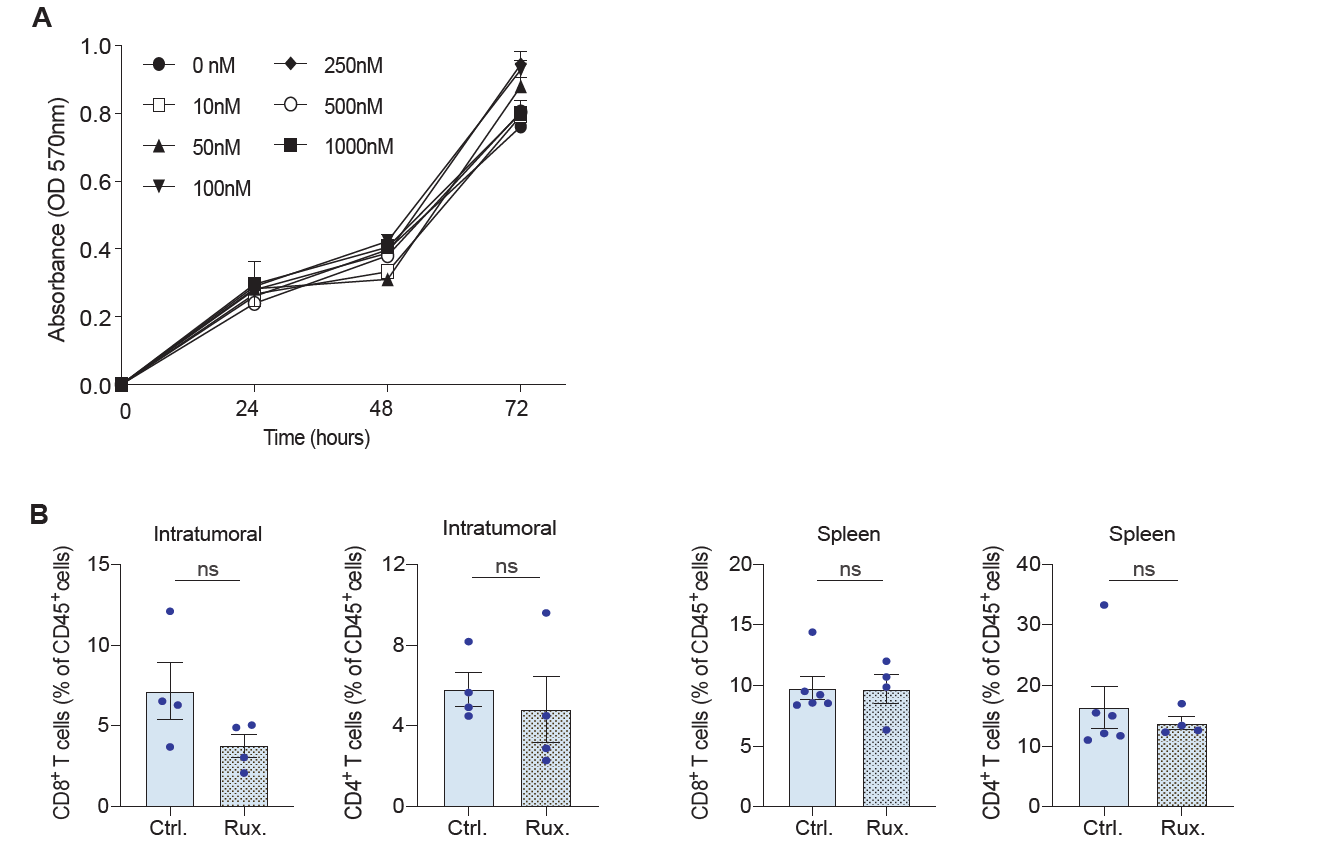

**Supplementary Figure 13. Ruxolitinib affects neither mT3-2D cellular proliferation rate *in vitro* nor T cell frequency and activity *in vivo*. (A)** Crystal violet proliferation curves for mT3-2D cells *in vitro* treated for 3 days with a range of ruxolitinib concentrations. **(B)** Flow cytometry analysis of T cells in mice treated with PBS (control) or ruxolitinib. Data represented as mean (±SEM). Mann-Whitney test was used. (rux.= ruxolitinib, ctrl.= control, ns=not significant).
